# The Malaria Cell Atlas: a comprehensive reference of single parasite transcriptomes across the complete *Plasmodium* life cycle

**DOI:** 10.1101/527556

**Authors:** Virginia M. Howick, Andrew J. C. Russell, Tallulah Andrews, Haynes Heaton, Adam J. Reid, Kedar Natarajan, Hellen Butungi, Tom Metcalf, Lisa H. Verzier, Julian C. Rayner, Matthew Berriman, Jeremy K. Herren, Oliver Billker, Martin Hemberg, Arthur M. Talman, Mara K.N. Lawniczak

## Abstract

Malaria parasites adopt a remarkable variety of morphological life stages as they transition through multiple mammalian host and mosquito vector environments. Here we profile the single-cell transcriptomes of thousands of individual parasites, deriving the first high-resolution transcriptional atlas of the entire *Plasmodium berghei* life cycle. We then use our atlas to precisely define developmental stages of single cells from three different human malaria parasite species, including parasites isolated directly from infected individuals. The Malaria Cell Atlas provides both a comprehensive view of gene usage in a complex eukaryotic parasite and an open access reference data set for the study of malaria parasites.

**One Sentence Summary:** Single-cell transcriptomics of malaria parasites illuminates gene usage across the entire life cycle, and allows precise developmental time assignment of parasite cells from distantly related *Plasmodium* species.

---

Single-cell RNA-sequencing (scRNA-seq) is revolutionising our understanding of heterogeneous cell populations, revealing rare cell types, unraveling developmental processes, and enabling greater resolution of gene expression patterns than has previously been possible (*1*). The ambition of cataloging the complete cellular composition of an animal is already becoming reality (*2, 3*), but thus far, atlasing efforts have focused on multicellular organisms. Here, we present the first comprehensive cell atlas of a unicellular eukaryote, the malaria parasite, across the entirety of its life cycle.

Although malaria parasites are unicellular, they display remarkable cellular plasticity during their complex life cycle with stages ranging from 1.2 to 50 μm and spanning vastly different human and mosquito environments. Clinical symptoms of malaria result from asexual replication within red blood cells, while transmission to new hosts relies on replication in the mosquito. Both disease development and transmission are therefore underpinned by the parasite’s ability to serially differentiate into morphologically distinct forms, including invasive, replicative, and sexual stages (Fig. 1A). This versatility is orchestrated by tight regulation of a compact genome, where the function of ~40% of genes remains unknown (*4*). Better understanding of gene use and gene function throughout the parasite’s life cycle is needed to inform the development of much-needed new drugs, vaccines, and transmission blocking strategies.

To begin to build the Malaria Cell Atlas, we profiled 1787 single-cell transcriptomes across the entire life cycle of *Plasmodium berghei* using a modified Smart-seq2 approach (*5*). Purification methods were adapted to isolate each stage of the life cycle, including challenging samples such as rings, which have low levels of RNA, and ookinetes, which are difficult to sort (fig. S1). Ninety percent of sequenced cells passed quality control (1787/1982 cells) and poor quality cells were identified per stage based on the distribution of the number of genes per cell (fig. S2). After quality control, we detected a mean of 1527 genes per cell across the entire data set; however, the number of genes detected was highly dependent on parasite stage (*p*<0.001; fig. S2). Transcriptomes were normalized with Trimmed Mean of M-values (TMM) in groups of related stages for further analysis. For samples expected to be overlapping or heterogeneous (*e.g*., the blood stages), we used k-means clustering to delineate stages and confirm their classification based on known marker genes and correlations with bulk reference data sets (fig. S3, S4). This allowed for differentiation between male, female, and asexual stages in the blood, as well as between ookinetes and oocysts in the heterogeneous population of parasites taken from the mosquito midgut in which ookinetes were actively invading.

All cell transcriptomes were visualised using UMAP (*6*) (Fig. 1B) and the first three principal components (Fig. 1C). Strikingly, cells oriented along a developmental path and also to some extent grouped by cellular strategy and host environment (e.g., actively replicative stages such as trophozoites and oocysts are near each other in UMAP and PCA, while PC3 separates the cells by host; Fig. 1B, 1C). All stages displayed marker genes concordant with known expression patterns (fig. S5). Merozoites, rings, trophozoites and schizonts formed a circle capturing the cyclical nature of the asexual intra-erythryocytic developmental cycle (IDC); (Fig. 1B, fig. S3). For 44 h liver schizonts, we also captured the host’s transcriptome, confirming at a single cell level that the parasite’s developmental progression is independent of the host cell’s cell-cycle state (*7*) (fig. S6).

**Figure 1.**
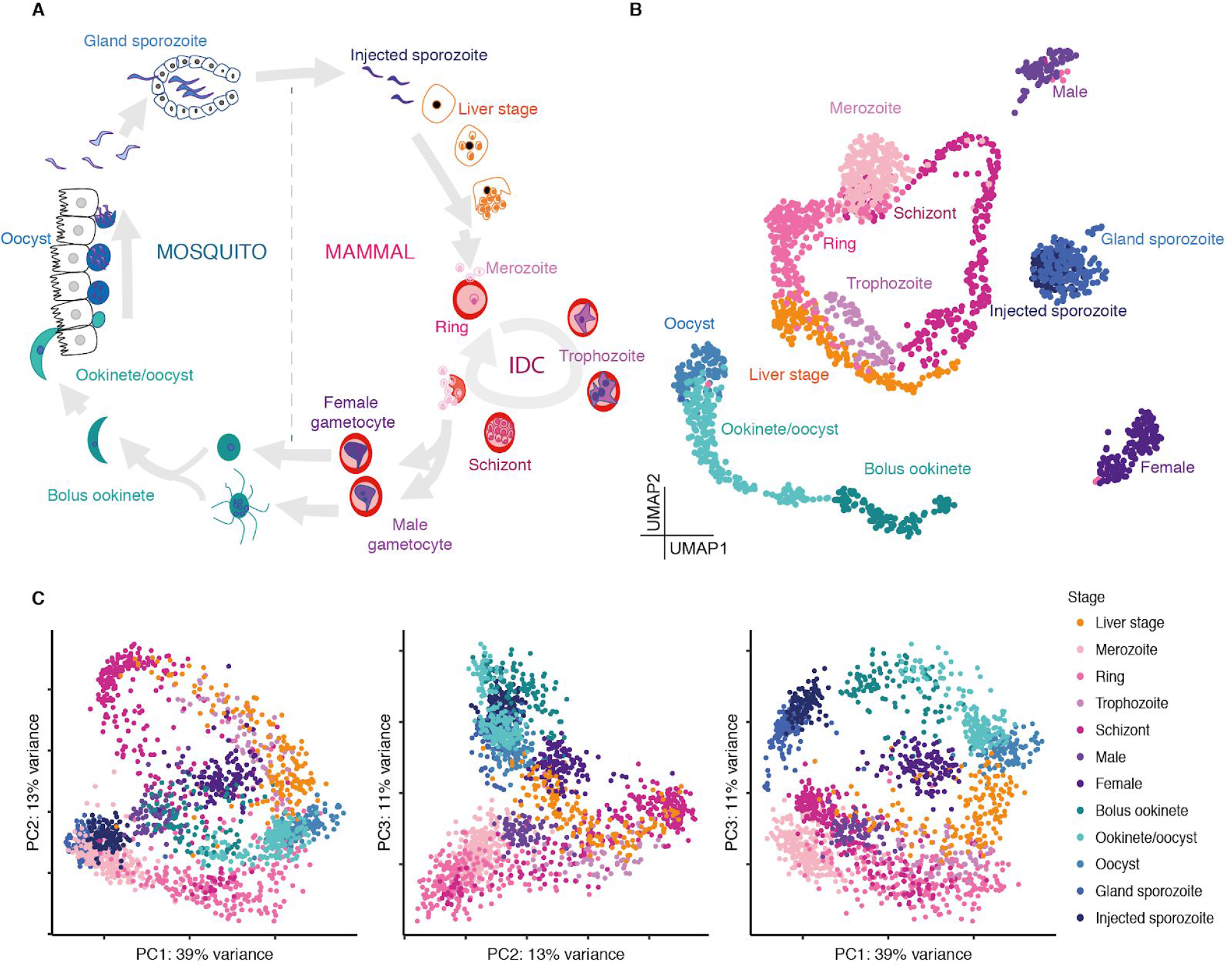
A single-cell atlas of the *P. berghei* life cycle. **(A)** The life cycle begins when an infected mosquito injects sporozoites into the mammalian host. From here, parasites enter the liver, where they develop, replicate, and then egress to enter the IDC. In the IDC, they invade erythrocytes, where they develop, replicate, burst, and re-invade erythrocytes cyclically. Sexual forms are taken up by the mosquito, and if fertilisation is successful, parasites invade the midgut and subsequently the salivary glands of the mosquito. In these different environments, parasites adopt different cellular strategies: replicative stages (liver schizont, blood stage schizont, oocyst), invasive stages (merozoite, ookinete, and sporozoite), and sexual stages (gametocytes). **(B)** UMAP of cells sampled from all stages of the life cycle, with cells colored according to their stage from (A). **(C)** The first three principal components from all stages in the life cycle.

Our survey of the *P. berghei* life cycle enables a global view of gene expression and prediction of function based on co-expression patterns. We constructed a k-nearest neighbour graph, where each node represents one of 5156 genes detected in the data set (file S1). Graph spectral clustering (*8*) was used to assign each gene to one of twenty modules based on the graph distance matrix (Fig. 2A, file S1). We observed gene clusters (1 and 2) consisting mainly of housekeeping genes and rRNA components that were highly expressed across the full life cycle. At the other extreme, some clusters (clusters 18-20) showed low overall expression and were primarily composed of genes from rapidly evolving multigene families (*pirs* and *fams*), which have no 1:1 orthologs with *Plasmodium falciparum* (fig. S7, Fig. 2B). Several gene clusters (clusters 7-16) were highly expressed in a single stage. We corroborated these stage-specific gene modules using two additional methods. First, we identified marker genes based on level of expression relative to all other stages (file S1). Additionally, for each stage we defined a core transcriptome of genes where transcripts were detected in >50% of cells (file S1). The number of genes unique to the core transcriptome for each canonical stage ranged from zero in merozoites to 237 in oocysts (file S1). Core genes for each stage were overrepresented in clusters that coincided with expression at that stage and contained genes involved in the cellular strategy of that stage (e.g. DNA replication, invasion, sexual development; fig. S8), confirming our module assignment to specific stages.

The majority of gene clusters show predominant expression in specific stages (Fig. 2B), offering new guidance as to where and how these genes might function. For example, CelTOS and CSP, both important invasion genes (*9, 10*), were found in cluster 16, which contains 79 genes most highly expressed in the invasive ookinete and sporozoite stages. Among these 79 genes are 34 annotated only as “conserved *Plasmodium* protein with no known function”. Their highly correlated expression with known invasion genes and their high expression in invasive stages will help inform future functional studies. We also overlaid asexual growth-rate data from a genome-wide knock-out screen (*11*). Genes expressed primarily in transmission stages (clusters 11-16) tended to show normal growth rates in asexual blood stages (Fig. 2C), offering further support that genes in these clusters are primarily important in transmission stages. For genes in each cluster, we also identified conserved motifs enriched in upstream regions as well putative binding sites for apetala2 (AP2) transcription factors that play critical roles in parasite progression through the life cycle (*12*) (fig. S9; file S1). This categorization will help to functionally annotate genes with unknown function, thereby enabling informed studies on gene regulation and supporting efforts to identify good candidates for transmission blocking drug and vaccine development.

**Figure 2-.**
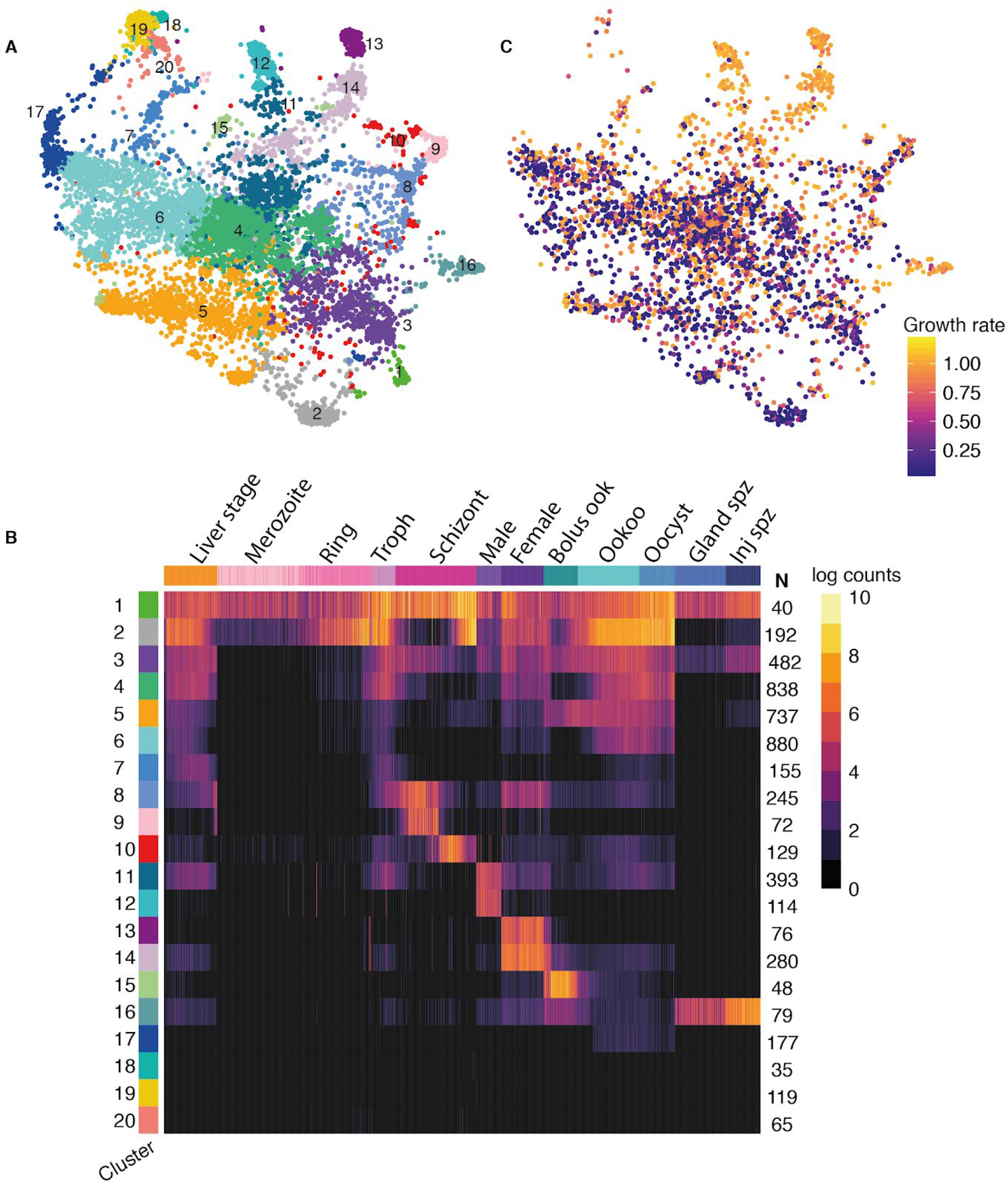
**(A)** A kNN force-directed graph of all 5156 detected genes. Each node represents a gene. Nodes are coloured according to their graph-based spectral clustering assignment. **(B)** A heatmap of mean expression for each cluster across all cells in the data set. Cells are ordered by their developmental progression. **(C)** The graph colored by relative growth rate phenotype of mutants in asexual blood-stage parasites (*11*).

Development is the primary driver of differences in gene expression across the life cycle. However, variation between individual parasites within developmental stages is important for adaptation to the host environment (*13*). The principal mechanism for intra-stage variation is thought to be driven by variation in expression amongst members of large multigene families whose functions are poorly defined (*14*). Most stages were enriched for variability in expression of such multigene families (fig. S10; file S1), the largest of which, *pir*, has a role in establishing chronic blood stage infections (*15*). Subsets of *pir* genes showed variable expression in different stages, coupled with distinctive upstream sequences (fig. S11). Such putative promoter architectures could define stage-specific expression, with epigenetic control defining which subset of members are expressed. Interestingly, we found five co-expressed pairs of *pir* genes in merozoites and rings (fig. S11). *Pir* genes within each pair were split across different chromosomes but shared similar promoter architectures, with different pairs having different promoters (fig. S11). While the function of these co-expression patterns is as yet unknown, such co-expression in a single cell can only be detected using scRNA-seq, highlighting another use of scRNA-seq towards identifying novel expression patterns.

Droplet-based approaches to generate single cell transcriptomes are nearly tenfold cheaper per cell than Smart-seq2, enabling the exploration of many more cells. In order to more deeply sample parasites along the entire IDC, we used the droplet-based 10X Chromium platform to simultaneously capture *P. berghei* and another parasite species, *P. knowlesi*, in a single inlet (*16*). We found that 6.34% of cells were dual-species doublets confirming a doublet rate as expected for conventional cells (fig. S12). After removal of doublets and additional quality control, we captured 4884 *P. berghei* cells and 4237 *P. knowlesi* cells.

We mapped *P. berghei* life stages between the 10X and Smart-seq2 technologies using Canonical Correlation Analysis (CCA) and scmap (Fig. 3A, fig. S12) (*17, 18*). CCA clustering shows good representation of all stages in both Smart-seq2 and 10X data sets within the IDC (fig. S12). Using scmap-cell, 94% of cells in the 10X data were assigned to a Smart-seq2 cell with high confidence, allowing us to align data sets (Fig. 3A). The additional *P. berghei* 10X data increases the coverage of cells in our atlas and confirms our ability to evaluate single cells characterised by different methodologies. To account for the continuous cyclical nature of the data, we ordered the 10X cells in pseudotime by fitting an ellipse to the first two principal components and calculating the angle around the centre of this ellipse for each cell relative to a start cell (Fig. 3B, Methods). Additionally, to confirm the orientation of the cycle with real time, we correlated each single-cell transcriptome with published bulk reference data sets and observed a high correspondence between bulk time point and pseudotime order (Fig. 3B, fig. S13).

We next used RNA velocity (*19*), which provides a snapshot of the dynamic state of each cell, to examine transcriptional rates across this deeply sampled set of *P. berghei* parasites. We find that transcription rates vary dramatically over the IDC, with peak transcription occurring in late rings, consistent with bulk studies of nascent RNA transcription (Fig. 3C, fig. S14) (*20*). Additionally, we generated a 10X data set comprising 6737 cells from the IDC stages of the human parasite, *P. falciparum*. We used scmap-cell to assign each *P. falciparum* and *P. knowlesi* cell to the *P. berghei* 10X reference index built with 1:1 orthologs, thus enabling us to align the developmental trajectories of these three species (Fig. 3C, fig. S14). We find the pace of transcription as measured by RNA velocity is similar across species, revealing conserved transcriptional rate in the intra-erythrocytic life of different malaria species in spite of vastly different hosts, and different life cycle lengths (24 hours for *P. berghei*, 27 hours for *P. knowlesi*, and 48 hours for *P. falciparum*; Fig. 3C, fig. S15).

Transcriptomic studies of both *in vivo* and *in vitro* malaria parasites are often confounded by multiple life stages within a single sample. The Malaria Cell Atlas can be used to deconvolve bulk transcriptomic data and identify the specific life stages that were present in a bulk RNA-seq sample. We demonstrate this using published data sets from *P. berghei* (*21, 22*) and *P. falciparum (23)* (fig. S16). Future bulk RNA-seq studies can use the atlas to identify differences in cell type compositions, potentially regressing these out to calculate more accurate differential expression between conditions or samples.

**Figure 3.**
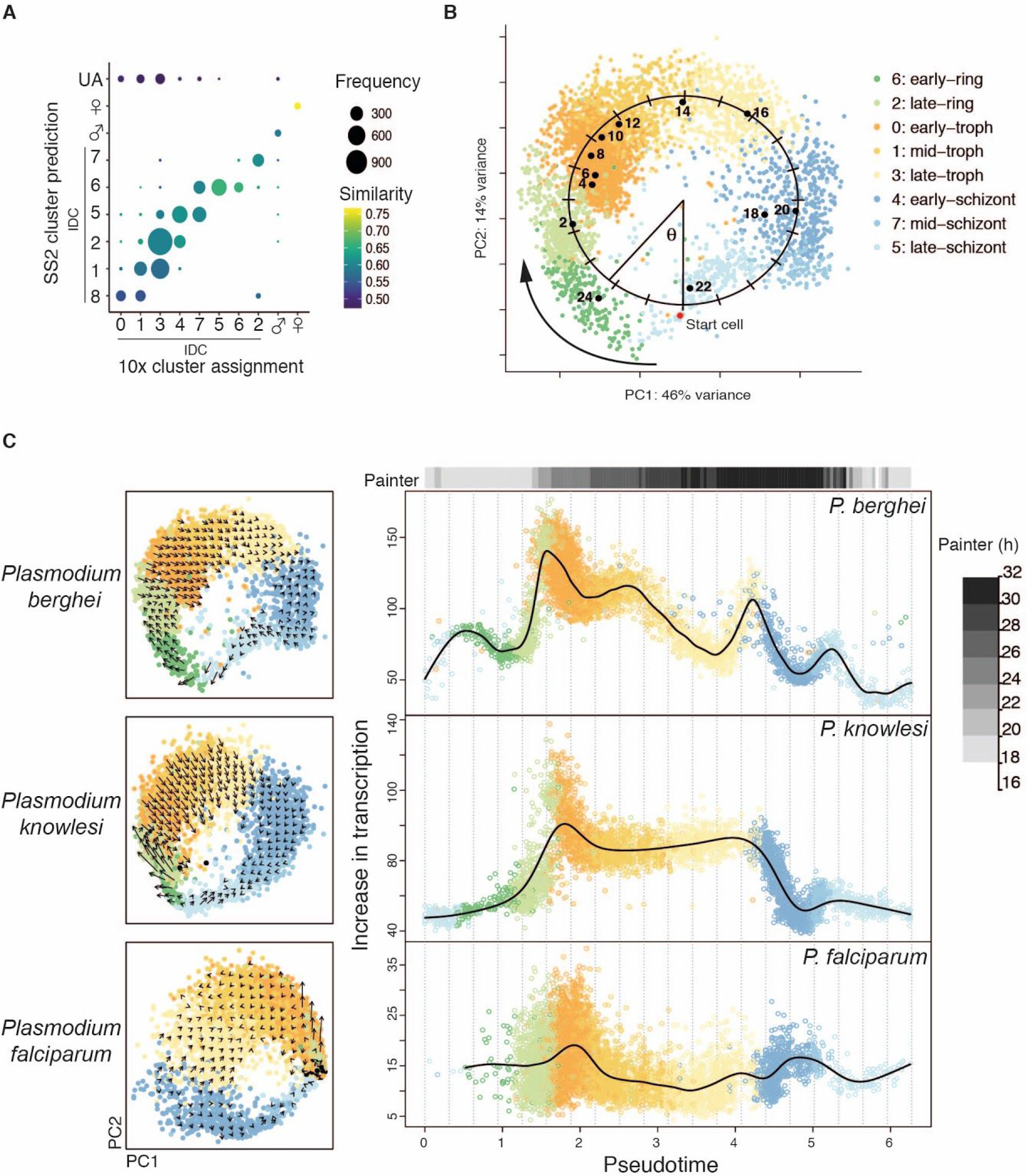
Alignment of data sets reveals conserved transcriptional rates in the IDC. **(A)** *P. berghei* 10X data mapped to Smart-seq2 data using scmap-cell. Cells are grouped according to their 10X cluster assignment (figs. S12, S14) and the SC3 (*24*) cluster of the Smart-seq2 cell it mapped to (fig. S14). 283 cells (< 6% of cells) were unassigned (UA) based on a cosine similarity threshold of 0.5. **(B)** PCA of *P. berghei* IDC cells from 10X. Pseudotime of each cell was measured by fitting an ellipse to the data and calculating the angle (radians) around the centre of this ellipse for each cell relative to the start cell (red point). Black points represent the mean PCA coordinates of the bulk prediction for each cell (*21*)(fig. S13). **(C)** Left: PCAs of three *Plasmodium* species colored by their *P. berghei* cell assignment based on scmap. Arrows represent the relative change in transcriptional state based on RNA velocity. Right: the scaled increase in expression over the IDC. Cells are ordered based on the pseudotime of their scmap assigned cell in the IDC *P. berghei* index. All three species show an abrupt increase in transcription at the same equivalent pseudotime early in the IDC (ring), followed by a steady decrease in later stages. The top bar represents the matched time point between the *P. berghei* RNA velocity-derived transcription rates and *P. falciparum* transcription rates reported by Painter et al. (*20*) using Pearson’s correlations.

*In vitro* systems, while critical for experimental studies of *Plasmodium* parasites, are unlikely to fully capture the breadth of expression variation of parasites circulating in naturally infected carriers. Moreover, several human-infecting species do not have any existing expression data and cannot be cultured *in vitro* (Fig. 4A). We therefore explored whether scRNA-seq on wild parasites taken straight from infected people could be placed in developmental time using our atlas. We developed a methanol-based preservation protocol that produced Smart-seq2 transcriptomes of equivalent quality to unpreserved cells in the lab (fig. S17). Next, we used the protocol to preserve samples from three naturally infected asymptomatic carriers in Mbita, Kenya, which we then sorted and sequenced in the UK. We recovered single cell transcriptomes from all three volunteers, and these field collected samples displayed similar quality to laboratory samples (fig. S17). *P. falciparum* cells mapped to our atlas revealed male, female, and early asexual parasites (Fig. 4B), which are the expected circulating stages for this species (Fig. 4A). Cells clustered by stage and not by donor indicating that comparisons both within host and between host are possible, and scRNA-seq on field parasites will enable transcriptional characterisation of natural infections. One of the volunteers was also infected with *P. malariae* leading to the first transcriptomic data for this species. Notably, we observed late developmental stages, which is expected as unlike *P. falciparum*, *P. malariae* late stages do not sequester in the deep tissue (Fig. 4). As a proof of concept, we have shown that parasite species that have previously been inaccessible for expression analysis can now be characterised by combining scRNA-seq with the atlas.

The Malaria Cell Atlas reference data set catalogs transcriptomes of every life stage along the parasite’s life cycle at single-cell resolution, and spans different technologies and different parasite species. The data are freely accessible as a processed data set and user-friendly web interface (*25*). As such, this will be a key resource for the malaria community in the study of transcriptional regulation and control of developmental progression at the highest resolution. The Malaria Cell Atlas provides a foundation for studying the biology of individual parasites directly from their natural environment, an important endeavor towards characterizing phenotypes critical for malaria control, including those related to pathogenicity, drug resistance, and transmission biology.

**Figure 4.**
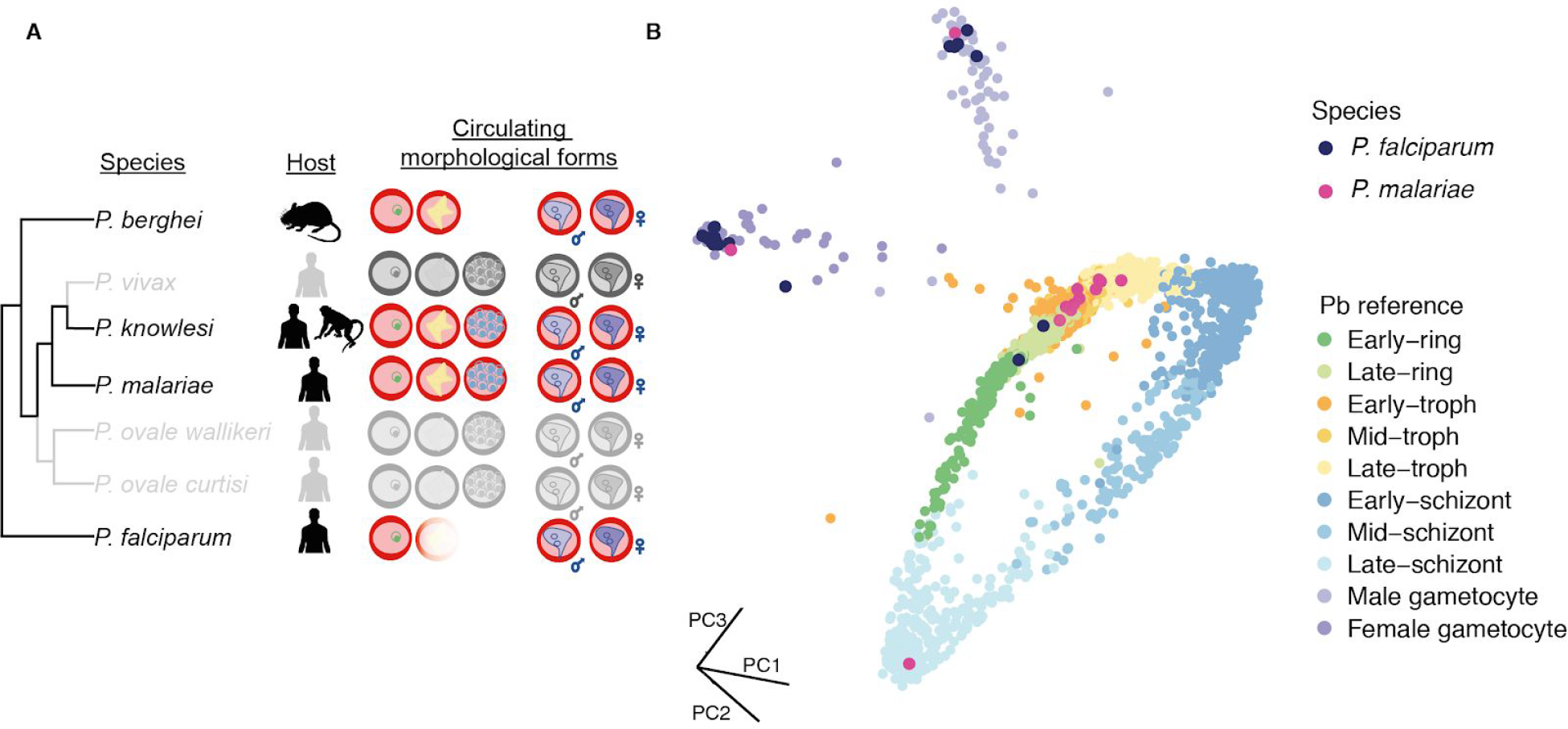
The Malaria Cell Atlas enables high-resolution mapping of field-derived single-cell transcriptomes of *P. falciparum* and *P. malariae*. **(A)** Phylogeny of *Plasmodium* showing the mammalian host and the stages found in circulation for each species. *P. falciparum* and *P. berghei* sequester their late stages in deep tissue, while other species have all morphological forms in circulation. Species in color were profiled in the atlas. **(B)** *P. falciparum* and *P. malariae* field-derived cells mapped onto the *P. berghei* 10X reference index using scmap-cell. The field-derived samples mapped to developmental stages that were expected in circulation for each species.

## Acknowledgements

The Wellcome Sanger Institute is funded by the Wellcome Trust (grant 206194/Z/17/Z), which supports MKNL, MH, OB, MB, and JR. MKNL is supported by an MRC Career Development Award (G1100339). KNN received funding from Danish institute of Advanced Study (D-IAS) and University of Southern Denmark (SDU). JKH and HB were supported by the Wellcome Trust (107372), the UK Department for International Development (DFID), the Swedish International Development Cooperation Agency (Sida), the Swiss Agency for Development and Cooperation (SDC), the German Academic Exchange Service (DAAD) and the Kenyan government. JR and LV were supported by NIH/NIAID R01AI137154. The authors would like to thank the staff of the Illumina Bespoke Sequencing and Core Cytometry teams at the Wellcome Sanger Institute for their contribution.

## Data and materials availability

All raw sequencing data are on ENA (ERP110344), and processed data are accessible via the Malaria Cell Atlas website (*25*). Analysis code is available at https://github.com/vhowick/MalariaCellAtlas.

## Supplementary Materials

### Materials and Methods

#### Parasite culturing *in vivo* and *in vitro*

*P. berghei* parasites came from drug selection marker-free reporter line (RMgm-928) that expressed mCherry, under the control of the *hsp70* promoter, throughout the life cycle (*26*). Parasites were propagated in female 6- to 8-week-old Theiler’s Original outbred mice supplied by Envigo UK. Mosquito infections were performed in two to five day old *Anopheles stephensi* mosquitoes.

*P. falciparum* (3D7) was maintained in O+ blood using RPMI 1640 culture medium (GIBCO) supplemented with 25 mM HEPES (SIGMA), 10 mM D-Glucose (SIGMA), 50 mg/L hypoxanthine (SIGMA), 10% human serum (obtained locally in accordance with ethically approved protocols), in a mix containing 5% O_2_, 5% CO_2_ and 90% N_2_.

*P. knowlesi* (strain A1-H.1) was maintained in continuous culture (*27*) in O+ blood, using RPMI 1640 culture medium (GIBCO) supplemented with 25 mM HEPES (SIGMA), 22.2 mM D-Glucose (SIGMA), 50 mg/L hypoxanthine (SIGMA), 0.5% (wt/vol) Albumax II and 10% horse serum, in a mix containing 5% O_2_, 5% CO_2_ and 90% N_2_. Cultures were maintained for >6 weeks without synchronisation to ensure good representation of all stages in the IDC.

Human O+ erythrocytes were supplied by NHS Blood and Transplant, Cambridge, UK. All samples were anonymized. Use of erythrocytes from human donors for *Plasmodium* culture was approved by the NHS Cambridgeshire 4 Research Ethics Committee (REC reference 15/EE/0253) and the Wellcome Sanger Institute Human Materials and Data Management Committee.

#### Parasite isolation, cell sorting and library preparation for Smart-seq2 scRNA-seq

##### Isolation of extraerythrocytic forms from HeLa cells

HeLa cells were cultured in DMEM supplemented with 10% FCS. *P. berghei* sporozoites were produced by homogenisation of 50 dissected sets of salivary glands from female *An. stephensi* mosquitoes 22 days after an infectious blood meal. Sporozoites were counted on a hemocytometer, resuspended in DMEM and added to an 80% confluent monolayer of HeLa cells at multiplicity of infection of one. The plate was spun at 300 g for 3 min and incubated at 37°C for 2 h, cells were then washed twice with PBS and placed back in complete medium. After 24 h cells were split back at 70% confluency. Cells were harvested by trypsinization 44 h after infection, washed once in PBS and sorted immediately.

##### Isolation of blood-stage merozoites

*P. berghei* parasites were purified from an overnight (24 h) 50 mL culture with 1 mL of infected blood using a 55% Histodenz cushion (SIGMA) as detailed elsewhere (*28*). Purified schizont stages were stained with Hoechst 33342 (ThermoFisher) at a final concentration of 2.5 μg/ml for 10 min on ice, pelleted at 450 g for 3 min, resuspended in 1 mL of medium and passed through a 1.2 μm filter (Pall Life Sciences). Merozoites in the filtered fraction were sorted immediately.

##### Isolation of ring-stage parasites

A mouse infected with RMgm-928 was terminally bled by cardiac puncture using a syringe containing heparin. The ~1 mL blood sample was immediately transferred onto ice and stained with 2.5 μg/ml Hoechst 33342 in PBS for 15 min (along with unstained controls for cell sorting). Cells were washed for 3 min at 800 g in RPMI and then once more for 3 min at 800 g in PBS. Parasites were then incubated in 0.02% saponin for 3 min, and then spun down at 1100 g at 4°C. Parasites were washed once in PBS for 3 min at 1100 g and then resuspended in 1 mL PBS for FACS.

##### Isolation of ookinetes from the blood bolus

Ookinetes were isolated from the blood bolus of *An. stephensi* midguts at 18 and 24 h post blood feeding from an RMgm-928 infected mouse at approximately 5% parasitemia. A lateral incision was made along the dissected mosquito midgut tissue to release the blood bolus and remaining blood was rinsed out using a syringe with PBS. Boluses from five midguts were pooled, diluted in 500 μl PBS and stained with SYBR green. In order to discriminate ookinetes from other stages in the blood bolus, a control feed was performed using a HAP2^−^-mCherry infected mouse. HAP2 is essential for fertilization, so the bolus contained parasites but no ookinetes (*29*). This allowed us to enrich our sample for ookinetes by gating on the level of mCherry and SYBR green fluorescence (fig. S1).

##### Isolation of invading ookinetes and oocysts from the midgut

At 48 h and four days post-bloodmeal invading ookinetes and oocysts were isolated from ten pooled infected midguts. Dissected midguts were disassociated in 200 μl of an enzymatic cocktail of collagenase IV (1 mg/ml) and elastase (1 mg/ml). The dissociation mixture was incubated at 30°C for 30 min with shaking at 300 rpm. Every 15 min tissue was mechanically disrupted by pipetting up and down 40 times. In order to capture only invading ookinetes at 48 h, the remaining blood bolus was removed as described above. As a control, midguts from mosquitoes that had fed on a HAP2^−^-mCherry mouse were disassociated to confirm that no remnants of the blood meal and non-invading parasites remained in the gut.

##### Isolation of salivary gland and injected sporozoites

Salivary glands from 20 *An. stephensi* infected with RMgm-928 were dissected on day 26 post-bloodmeal. Sporozoites were released from the glands by homogenising the samples manually with a pestle in PBS. Samples were filtered with a 20 μm filter prior to sorting to remove large fragments of mosquito tissue. Simultaneously, female *An. stephensi* mosquitoes from the same infectious feed were fed using a standard membrane feeding assay containing approximately 600 μl fructose solution (80 g/L) with 10% human serum (filter sterilized and heat inactivated). Mosquitoes were exposed to the feeder for 12 min. After this, the remaining fructose/serum solution was removed from the feeder and the presence of sporozoites in this solution was microscopically confirmed. Samples were then taken directly to cell sorting.

##### Preservation and isolation of cells from fresh peripheral blood samples

Samples were procured in the district of Mbita (Kenya) from asymptomatic volunteers in accordance with a study protocol reviewed and approved by the KEMRI Scientific and Ethics Review unit (KEMRI/RES/7/3/1). Following screening with a rapid diagnostic test (SD BIOLINE™ Malaria Ag P.f/Pan (HRP-II/pLDH)), venous blood samples from infected volunteers was collected in EDTA-vacutainers. 1 mL of each sample was resuspended in 5 mL of suspended animation buffer (10 mM Tris, 150 mM NaCl, 10 mM glucose, pH 7.37)(*30*) and placed on a magnetic column (MACs, Miltenyi). Late stage parasites were eluted, washed once in suspended animation buffer and resuspended in 200 μl of PBS. Samples were then fixed with 800 μl of methanol (Sigma) and preserved at −20°C. Another 1 mL was leucodepleted with a Plasmodipur filter (EuroProxima), washed twice in PBS, lysed twice with 0.15% Saponin (Sigma), washed twice in PBS and resuspended in 200 μl of PBS. Samples were then fixed with 800 μl of methanol (Sigma) and preserved at −20°C. Samples were rehydrated with PBS and stained with 2.5 μg/ml Hoechst in PBS for 15 min, and washed once in PBS prior to sorting.

##### Cell sorting

All parasite cell sorting was conducted on an Influx cell sorter (BD Biosciences) with a 70 μm nozzle. The HeLa samples were sorted on a Sony SH800 with a 100 μm nozzle chip. Parasites were sorted by gating on single cell events and mCherry fluorescence (all stages) or Hoechst fluorescence (merozoites, field parasites). All parasites were sorted into nuclease-free 96 or 384 well plates (ThermoFisher) containing lysis buffer as described previously (*5*). Sorted plates were spun at 1000 g for 10 seconds and immediately placed on dry ice.

##### Library preparation and sequencing

Reverse transcription, PCR, and library preparation were performed as detailed previously (*5*). All libraries were prepared in 96-well plates except a single 384-well plate of late blood stages. In the latter case the lysis buffer volume was reduced to 2 μl, and the elongation temperature of the PCR was reduced to 68°C. Cells were multiplexed up to 384 and sequenced on a single lane of HiSeq 4000 with 75 bp paired-end reads.

#### Parasite preparation and loading of 10X scRNA-seq

##### Parasite preparation

For *P. berghei* samples, blood was obtained by terminal bleed and passed through a pre-wetted Plasmodipur syringe filter (Europroxima) to filter out white blood cells prior to culturing. Three cultures were generated: cultured for 30 min, 10 h, and 20 h at 36.5°C with shaking at 65 RPM. Cultures were smeared prior to harvesting in order to ascertain parasitemia. After harvesting, the total number of red blood cells in each sample was counted using a disposable hemocytometer. This count was corroborated using a Countess cell counter. The number of infected red blood cells in each culture was used as a cell count and cells were pooled 1:1:1 from the three time points and kept on ice. For *P. knowlesi*, the parasitemia of the cultured desynchronized parasites was measured and then cells were harvested by centrifugation at 450 g for 3 min at 4°C. Supernatant was removed and parasites were washed twice in PBS before resuspension in PBS. The concentration of red blood cells was then calculated by manual hemocytometer, before calculating the final infected red blood cell concentration using the parasitemia. Cells were then pooled 1:1 with the *P. berghei* cell mixture described above in order to run a dual species 10X analysis to evaluate doublet rates. *P. falciparum* parasites were prepared in the same manner as *P. knowlesi* but were run on their own 10X inlet.

##### 10X loading

Cells were loaded according to manufacturer’s instructions to recover 10000 cells per inlet. Chromium 10X v2 chemistry was used and libraries were prepared according to manufacturer’s instructions. Each 10X input library was sequenced across two Hiseq 2500 Rapid Run lanes using 75 bp paired-end sequencing.

#### Bulk transcriptomics

Three *P. berghei* samples were prepared for bulk RNA-seq including early asexuals, late asexuals, and ookinetes. Mice infected with hsp70p:mCherry *P. berghei* were terminally bled by cardiac puncture using a syringe containing heparin. For the two asexual samples, the blood was treated with ammonium chloride to remove uninfected erythrocytes (*31*) either straight after the bleed (early) or after 24 h of *ex vivo* culture (late). For the ookinete sample, the blood was cultured for 24 h, as described (*32*). RNA was extracted with TriZol according to the manufacturer’s recommendations, assayed with an Agilent RNA 6000 Nano assay and transcriptomes were generated as described. A modified RNA-seq protocol was used. PolyA+ RNA (mRNA) was selected using magnetic oligo-d(T) beads. Reverse transcription using Superscript III (Life) was primed using oligo d(T) primers; second strand cDNA synthesis included dUTP. The resulting cDNA was fragmented using a Covaris AFA sonicator. A “with-bead” protocol was used for dA-tailing, end repair and adapter ligation using “PCR-free” barcoded sequencing adaptors (NEB) (*33*). After two rounds of SPRI cleanup (Agencourt) the libraries were eluted in EB buffer and USER enzyme mix (NEB) was used to digest the second strand cDNA, generating directional libraries. The libraries were quantified by qPCR and sequenced on an Illumina HiSeq 4000.

#### Mapping and generation of expression matrices for scRNA-seq transcriptomes

##### Smart-seq2 mapping

Single-cell *Plasmodium* transcriptomes were mapped as reported previously (*5*). Briefly, trimmed reads were mapped using HISAT2 (v 2.0.0-beta) (*34*) to the *P. berghei* v3 genome (October 2016), and using STAR (v 2.5.0a) to the *P. falciparum* v3 (January 2016) and *P. malariae* v1 (March 2018) genomes using default parameters (*35*). Reads were summed against genes using HTseq (v 0.6.0) (*36*). For the co-expression of HeLa cells and liver stage parasites analysis, both HeLa cells and parasites were mapped to respective genomes with STAR (v 2.5.1b) using default parameters (*35*).

##### 10X data alignment, cell barcode assignment, and UMI counting

The Cell Ranger Single-Cell Software (version 2.1.0) was used to process sequencing reads, assigning each read to a cell barcode and UMI using standard parameters (*16*). After barcode assignment, the cDNA insert read was aligned using cell ranger (v 2.1.0) to a combined reference genome of *P. knowlesi* (March 2014) and *P. berghei* (July 2015), and the *P. falciparum* run was aligned to the 3D7 genome v3 (January 2016). These reference genomes were all obtained from https://www.sanger.ac.uk/resources/downloads/protozoa/.

#### Filtering and normalisation of scRNA-seq data

##### Smart-seq2 filtering and normalization

Poor quality cells were identified on a per stage basis based on the distribution of the number of genes per cell, given the high variability of genes detected between stages (fig. S2). Cells with fewer than 1000 genes per cell and 2500 reads per cell were removed from the liver stage parasites, trophozoites, male and female gametocytes, ookinetes, ookinetes/oocysts, and oocyst stages. Cells with fewer than 500 genes per cell and 2500 reads per cell were removed from schizonts and injected sporozoites. Cells with fewer than 40 genes per cell and 1000 reads per cell were removed from merozoites, rings, and gland sporozoites (fig. S2). Additionally, we removed genes from further analysis that were detected in fewer than two cells across the entire data set. The final data set contained 1787 high-quality single cells from 1982 sequenced and 5156 genes out of 5245 genes with annotated transcripts. Transcriptomes were normalized with the weighted Trimmed Mean of M-values (TMM) method (*37*). Cells were normalized either all together or in five groups containing biologically similar stages: groups included IDC, liver-stage, gametocytes, ookinetes/oocysts, and sporozoites. Visual inspection of the relative expression plot (fig. S2) showed little difference between normalization by biological group versus all together. Unless otherwise specified, further analysis was done on cells normalized by biological group.

##### 10X filtering and normalisation

For *P. berghei*, the output filtered matrix from Cell Ranger was read into Seurat (v 2.3.4) (*17*). Low quality *P. berghei* cells with fewer than 230 detected genes were removed from further analysis. Initial inspection of filtered cells in the *P. knowlesi* and *P. falciparum* data sets showed that early-stage and late-stage IDC parasites were missing. These stages express fewer genes per cell relative to later stages based on our Smart-seq2 data and we have previously observed a lower detection of genes per cell in *P. falciparum* (*5*), suggesting these cells may have been removed by Cell Ranger’s default thresholding. Using the raw output matrices for these species, we adjusted thresholds to retain cells with >100 genes per cell for *P. falciparum* and >150 genes per cell for *P. knowlesi*. Intraspecies doublets were identified and removed from all three species using doubletFinder (v 1.0.0) (*38*). For the *P. berghei/P.knowlesi* run, we identified interspecies doublets as cells that contained >50 UMIs that mapped to each species (1005 cells) (fig. S12A). The expected intraspecies doublet rate was calculated based on this interspecies doublet rate, the relative proportion of each species, and the additional quality control thresholding. For *P. falciparum*, the intraspecies doublet rate was calculated from the expected doublet rate table provided by 10X genomics. Thus, the number of intraspecies doublet cells removed were as follows: *P. berghei* = 200, *P. knowlesi* = 287, *P. falciparum* = 530 (fig. S12B). Doublets do not show a stage-specific bias (fig. S12B).

#### Single-cell transcriptome analysis of Smart-seq2 data

##### Cell clustering and projection

For timepoints where a heterogeneous population of stages was collected, we used k-means clustering using SC3 (version 1.7.7) to delineate stages and confirmed their classification based on known marker genes (*24*). This method was used for classification of males, females, trophozoites and schizonts, as well as ookinetes and oocysts (fig. S3, S4). For visualization in two dimensions we performed Uniform Manifold Approximation and Projection (UMAP) (*6*) with the python package umap version 0.1.1 using the correlation distance metric, k-nearest neighbors of ten, min_dist of one, spread of two, and bandwidth of one.

##### HeLa QC and cell-cycle analysis

We performed initial filtering to identify the most robustly expressed genes across single-cells. Genes were required to be expressed in >30 cells (of 164 cells) and cells needed to express >500 genes in both parasite and matched HeLa cells to be retained. This resulted in 163 matched cells with 4480 parasite genes and 8059 HeLa cell genes. We performed clustering of single HeLa and parasite cells independently using either all highly variable genes or subsets of annotated cell cycle genes. The highly variable genes are identified by plotting the averaged gene expression against gene dispersion (similar to SEURAT, (*17*)). Louvain clustering was performed on single HeLa cells using only cell cycle genes resulting in four louvain groups (fig. S6). These groups are highly indicative of cell cycle progression starting from group 0 (G0/G1) to group 1 (G1S) to group 3 (G2) to group 4 (G2M).

##### Pseudotime

To order cells in a developmental trajectory, we reconstructed pseudotime using SLICER (*39*). Variable genes were identified within SLICER and then selected to build the trajectory based on a neighbourhood variance that identifies genes that vary smoothly across the cell sets. SLICER was run independently on three groups of cells: (1) the liver stage parasites, (2) the entire IDC (merozoites, rings, trophozoites and schizonts), (3) the ookinete to oocyst transition (bolus ookinetes, ookinete/oocyst, and oocyst). We assessed the performance of the algorithm by confirming the pseudotime order matched the ground truth time point collections and expression of known marker genes over development (*e.g.* fig. S4). To order all cells across the life cycle (Fig. 3B, fig. S10, S11), we compiled these pseudotime orders with known timing of other stages that did not show a developmental signature (mature gametocytes and sporozoites).

##### Gene clustering and visualization of Smart-seq2 data

The gene count matrix was normalized by dividing by the mean counts for each gene and log scaling. This was done to reduce the amount to which gene clusters were driven by total gene expression and instead focus on the pattern of expression across cells. A k-nearest neighbor (kNN) graph was formed on the gene normalized expression matrix with the Nearest Neighbors subpackage of python’s scikitlearn version 0.19.2 with parameters of k = 5 and a manhattan distance metric (*40*). k of 5 was chosen because it was smaller than the smallest cluster we were interested in detecting and the graph appeared robust from k=3 to k=20. We then performed spectral graph clustering on this kNN graph using the SpectralClustering subpackage of python’s scikitlearn version 0.19.2 (*8, 41*). The graph was visualized in Gephi with the forceatlas 2 graph layout algorithm in linlog mode to better show the clustering structure of the data (*42, 43*).

##### Marker genes

Marker genes for each stage were identified in two ways. Firstly, differentially expressed genes were calculated using the findMarkers function in scran (*44*). This function performs a Welch t-test between pairs of stages and then identifies genes that are uniquely expressed in that cluster (pval.type = “all”, direction = “up”). This method was used to identify markers for each canonical stage, as well as marker genes within each host (mouse vs. mosquito) and each cellular strategy (invasive, replicative, and sexual forms) (file S1). Secondly, marker genes were identified by defining a core set of genes for each stage as all genes that are expressed in more than fifty percent of cells. To avoid bias from the number of cells sampled within a stage, sixty cells were randomly selected per stage. We defined the unique core transcriptome as genes from each stage’s core that were unique to that stage’s core (i.e., not also found in more than 50% of cells from any other stage) (file S1).

##### Motif Discovery

Motif discovery was performed using DREME which searches for short (8 bp) motifs expressed as regular expressions (consensus sequences allowing for wildcards but not variable length gaps) in a given set of sequences (*45*). The 1000 bp upstream of the start codon for each gene detected in the Smart-seq2 data set was used in the analysis. For each cluster the input data set was the upstream regions of each gene within that cluster, and the negative set was the upstream region of genes that were not in that cluster.

##### Analysis of development-independent gene expression variability

Almost all genes in *Plasmodium* genomes vary in expression over the life cycle. This is mainly thought to be related to development of the parasite as it transitions between different life stages. We first identified highly variable genes in each stage independently. In the Smart-seq2 data, we used a general linear model to regress out the effect of pseudotime within developing stages (liver stage exo-erythrocytic forms, or EEFs, merozoites, rings, trophozoites, schizonts, ookinetes and oocysts). We preserved the mean expression of each gene by adding the predicted value of the mean to the residuals of the general linear model, in addition we set any negative corrected values to zero in order to preserve the non-negativity of gene expression values. Finally, since the correction often shifted zeros to values only slightly above zero, we rounded these values down in order to meet the assumptions of the M3Drop model. We then used M3Drop (*46*) to identify genes with remaining heterogeneity (False Discovery Rate < = 0.05), adjusted for mean-expression level. Enrichment of each gene family within each stage was determined using the hypergeometric test with correction by FDR and a cut off of 0.05.

We examined *pir* gene promoter architectures to determine whether particular gene expression patterns might be driven by transcription factors. Firstly, we identified the 5’ UTR and upstream intergenic (putative promoter) regions of the *pir* genes shown in fig. S11. This was done manually by browsing the genome and referring to three *P. berghei* bulk RNA-seq samples of mixed early and late asexual stages as well as ookinete stages. Illumina reads from these libraries were mapped to the *P. berghei* v3 genome sequence using HISAT2 v2.0.0 (*34*), with --rna-strandness RF --max-intronlen 5000. The data were viewed using Artemis v18.0.0 (*47*). 5’ UTRs were defined as the region between the start codon and where RNA-seq coverage dropped to zero in at least two of the three samples. Upstream intergenic regions were defined from the start of the 5’ UTR to the next, upstream increase in coverage from one or more RNA-seq libraries. The upstream intergenic regions were BLASTed against each other (blastall 2.2.25, −p blastn −e 1e-20). The sequences involved in each hit were extracted, excluding those overlapping others with lower E-values. These sequences were then BLASTed against each other (blastall 2.2.25, −p blastn −e 0.01) and the resulting similarity matrix was used to cluster them with MCL v12-068 (*48*) with the inflation parameter set to 1.4. Sequences were collected together based on the clustering and aligned using MUSCLE v3.8.31 (*49*). The alignments were then trimmed by identifying highly conserved regions. Alignments in non-overlapping windows of ten nucleotides were evaluated, counting the proportion of sequences that were ungapped. An alignment position was called as good if >= 70% of sequences were ungapped at that position. A window of ten nucleotides was called as a block if it contained no more than three bad positions. If there was more than one bad block in a row, a conserved region was ended. Only the longest conserved region from an alignment was kept. Sequences which began or ended within the conserved region were then removed. These alignments were used to build nucleotide profile Hidden Markov Models (HMMs) using HMMer i1.1rc3 (*50*). The models were then searched against the *P. berghei* v3 genome sequence, also using HMMer, to identify further members of the sequence families. Each hit was associated with the nearest downstream protein-coding gene. We identified eight upstream intergenic (promoter) sequence families associated with *pir* genes that we called A, C, D, F, G, H, I J (fig. S11)

##### RNA Velocity

For each IDC (ring, trophozoite, schizont) Smart-seq2 cell that passed quality control, the exonic, intronic, and mixed reads were counted using RNA velocity (*19*). Intronic and mixed reads were combined to estimate the total unspliced reads in the data, whereas purely exonic reads were assumed to represent spliced transcripts. Cells were split by life cycle stage as for the pseudotime analysis, and the expected ratio of spliced to unspliced reads for each gene was fit using RNA velocity, and residuals for each cell were estimated for each group independently. To ensure we only considered genes that were fit well by the RNA velocity model, we required a minimum slope of 0.1 (increased from the default setting of 0.05) and a minimum correlation between spliced and unspliced reads of 0.5 (increased from the default setting of 0.05). To improve fits, we used the cells with the top and bottom 7.5% of expression levels for the fitting. Genes where over 90% of residuals were either positive or negative were excluded as poorly fit genes. This resulted in 1,345 genes × 548 cells for the IDC.

#### 10X single-cell transcriptome analysis

##### Cell clustering

To identify male and female gametocytes in the *P. berghei* data, data were log normalized, and clusters were identified using the shared nearest neighbor modularity optimization based clustering algorithm in the FindClusters() function in Seurat (*17*). Two clusters corresponded to gametocytes based on expression of marker genes. These clusters were removed when comparing the data to the Smart-seq2 via CCA, as well as for the pseudotime assignment, and alignment of the three data sets in scmap (Fig. 3). The three species IDC PCAs were generated on TMM normalized data in scater (version 1.6.3) (*51*). Additionally, we identified clusters using the CCA in Seurat to compare the two methods of scRNA-seq (Smart-seq2 and 10X) (*17*). We identified nine clusters of cells that had good representation in both data sets (fig. S12C). One cluster, “8”, contained only 15 cells across the two data sets and was removed from further analyses.

##### SCmap

We used scmap (version 1.1.5) (*18*) to compare data sets. We built three sets of cell indices that could be queried with the scmapCell() function that would allow each individual cell in the query data set to be mapped to a reference index (*18*). To compare the Smart-seq2 and 10X data we built an index of the blood-stage Smart-seq2 data (including gametocytes) and mapped the full *P. berghei* 10X data set (including gametocytes) onto it. Because the IDC consists of a continuous set of cell-stages and not discrete clusters, we modified the cell assignment method in scmap: cells were assigned based on the top nearest neighbor. If the top cell had a cosine similarity of greater than 0.5, the query cell was assigned to that indexed cell along with its supporting metadata (cluster assignment, bulk prediction, pseudotime value). Using this cosine similarity threshold, 94% of 10X *P. berghei* cells were assigned to a cell in the SS2 *P. berghei* reference data set.

To align the IDC trajectories across the three 10X datasets, we first compiled a set of one-to-one orthologs between ten *Plasmodium* species (*P. berghei*, *P. knowlesi*, *P. falciparum*, *P. malariae*, *P. ovale*, *P. vivax*, *P. gallinaceum*, *P. yoelii*, *P. chabaudi*, *P. cynomolgi*) from OrthoMCL (*52*) (file S1). Using these orthologs, we built an scmap reference index that contained all *P. berghei* 10X IDC cells (gametocytes removed). We mapped both the *P. falciparum* and *P. knowlesi* data to this ortholog reference index. In addition to identifying the top nearest neighbor cell, we were able to incorporate information from the top three nearest neighbors to assign each cell based on the principal component space. To do this we took a mean of the first two principal components of the top three nearest neighbors. Given this coordinate assignment, we located the nearest cell on the PCA and assigned the query cell to this index cell. If all three of the nearest neighbors had a cosine similarity of 0.3, then the query cell was given an assignment. With this lower cosine similarity threshold to account for cross species differences, we were able to assign over 96% of *P. falciparum* and 99% of *P. knowlesi* cells to a *P. berghei* index cell.

Finally, to map single cell samples from the field, we built a 1:1 ortholog index of the complete 10X *P. berghei* data set, including the gametocytes that were excluded for the IDC evaluations. We used this reference because (i) of its full representation of the IDC and mature gametocytes (ii) it originates from an *in vivo* system like the volunteer cells. Because the gametocyte data was more sparse, the IDC cell assignment was based on the top nearest neighbor alone along with a cosine similarity threshold of 0.4. Using this method, we were able map 13 *P. malariae* cells and 22 *P. falciparum* cells, assigning each cell to a developmental time.

##### ‘Clock’ pseudotime

For the three 10X data sets, pseudotime around the IDC was calculated by fitting an ellipse to the data projected into the first two principal components using direct least squares (Fig. 3C, fig. S14). Angles around the centre of this ellipse were calculated for each cell and oriented to a starting cell which was defined using known markers. In order to align the three species in pseudotime, 10X data from *P. knowlesi* and *P. falciparum* were projected directly onto the *P. berghei* reference using scmap (*18*) and cells were given the pseudotime of their *P. berghei* assigned cell. This ‘clock’ pseudotime was aligned to real-time progression through the IDC using two methods. First, bulk RNA-seq data (*21*), from synchronized *P. berghei* parasites across twelve equally spaced time points around the IDC was projected onto our single-cell reference and their position in the first two principal component space was estimated from the average of their three nearest neighbours. These principal component locations were used to calculate a respective pseudotime for each bulk sample. Secondly, we mapped our single-cell RNA-seq transcriptomes onto the densely sampled *P. falciparum* bulk RNA-seq time-course generated by Painter et al. (*20*). Genes were mapped across species using 1:1 orthologs (see above) and log-normalized RNA velocity-derived transcription rates were matched to the log-normalized transcription rates reported by Painter et al. using Pearson correlations.

##### RNA velocity

We also ran RNA velocity on the 10X single-cell RNA-seq data from each species independently. Cells were filtered as described above, genes were filtered to exclude genes that didn’t have at least one unspliced transcript in at least 10 cells and one spliced transcript in at least 20 cells. To account for the high number of zeros present in 10X data we increased the k for the cell and gene k-nearest neighbour smoothing included in RNA velocity to 50 and 5, respectively. To ensure good fits to the genes, we required a minimum slope of 0.2 and minimum correlation of 0.2 and used the top and bottom 20% of cells for the fitting. In addition, we excluded poorly fit genes as above. After this filtering, *P. knowlesi* data contained 1,235 genes × 4,237 cells, *P. berghei* data contained 1,368 genes × 4,763 cells, and *P. falciparum* data contained 645 genes × 6,737 cells.

##### Deconvolution of bulk transcriptomic samples using the scRNA-seq

The *P. berghei* 10X data was used as a reference and marker genes were called for each cluster in Seurat (v 2.3.4) using the Standard AUC classifier method Genes that were not detected in >40% of cells and negative markers were excluded. The top 10 marker genes in each cluster, by power, were identified and used for deconvolution (n = 107). BSeq-sc (v 1.0) (*53*) was then used to estimate the proportion of cell types in each bulk sample using the default analysis pipeline.

### Supplementary Figures

**Fig. S1.**
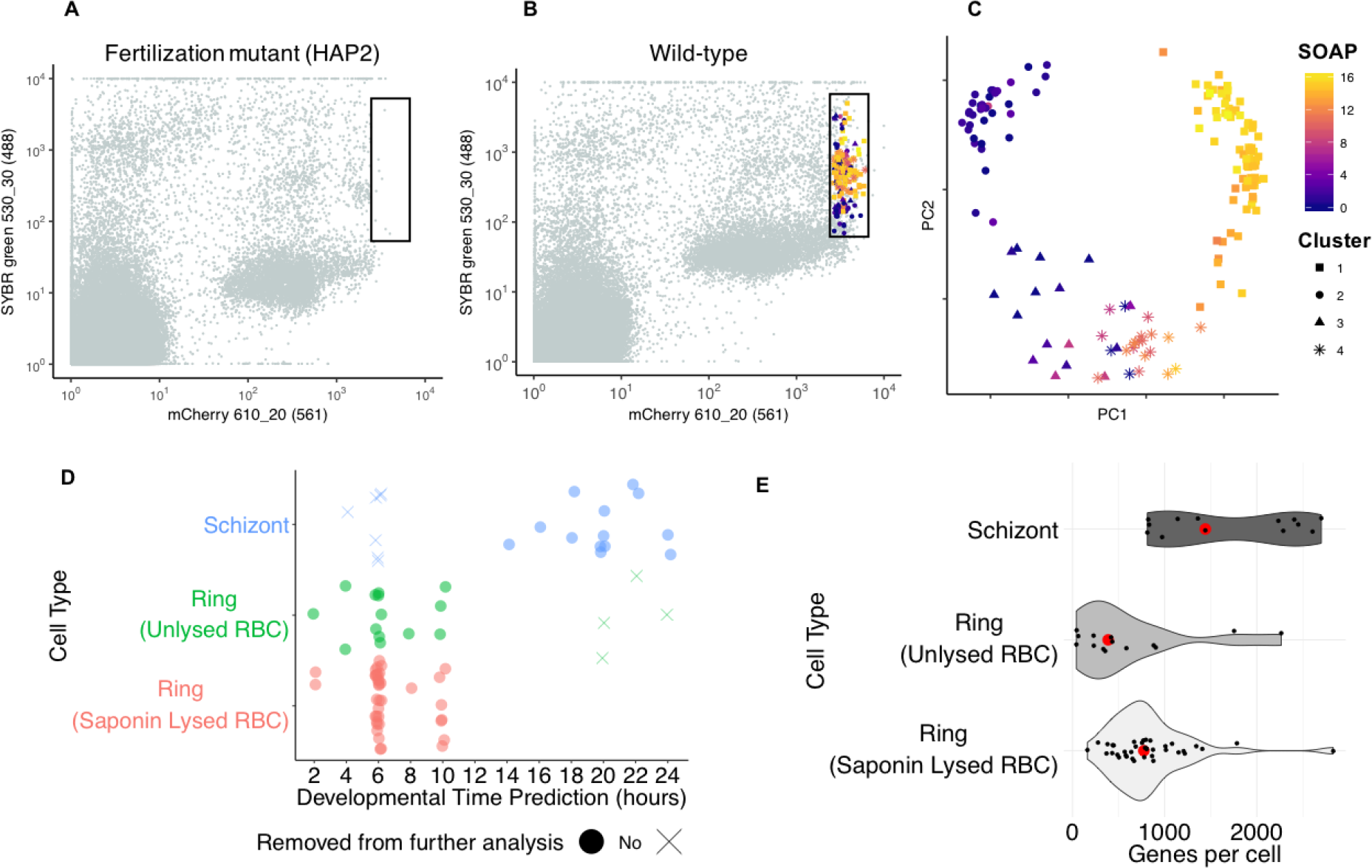
Isolation of ookinete and ring-stage parasites. **(A-C):** FACS gating for ookinete isolation from the blood bolus 18 and 24 h after feeding was determined using an mCherry expressing HAP2 mutant as a negative control. Parasites were expressing mCherry and stained with SYBR green. **(A)** FACS plot of the blood bolus contents from the HAP2 mutant control. **(B)** FACS plot of the wild type blood bolus. Cells that were sequenced are coloured by expression of the ookinete marker gene SOAP (*54*) and shaped by the SC3 cluster assignment. **(C)** A PCA of profiled cells coloured by expression of SOAP and shaped by SC3 cluster. Clusters 1 and 4 were maintained in the data set and likely represent mature ookinetes (cluster 1) and developing retorts (cluster 4). Cluster 3 may correspond to non-replicated zygotes: cells are expressing Pbs25 (*55*), but not SOAP. Cluster 2 may correspond to unfertilised parasites as there was inconsistent expression of canonical marker genes and generally lower levels of SYBR green. **(D-E):** To evaluate the protocol for saponin-mediated lysis of ring-stage infected red blood cells, a 96-well Smart-seq2 plate was sorted consisting of ¼ schizonts, ¼ unlysed rings, and ½ lysed rings. **(D)** The lysed and unlysed cells were collected from the same mouse controlling for life cycle stage. However, in order to confirm that we had sorted rings rather than early trophozoites, we used Spearman’s rank-order correlation to correlate each cell to published bulk time course data (*21*). The maximum r value for each cell was then chosen and the cell was assigned that time point. The jitter plot shows the predicted life cycle stage for each sorted sample, confirming similar predicted stages among lysed and unlysed rings. Data points are coloured according to the sorted population they belong to. Cells were then filtered according to their life cycle stage prediction, so as to remove cells that clearly did not belong to that cell type. Specifically, cells from the schizont culture that were predicted to be <10 h and and cells from the ring culture that were predicted to be >15 h were removed. Cells that were removed from further analysis are shown on the plot. **(E)** A violin plot showing the distribution of genes per cell per stage as well as the recovery rate for each stage. Lysing the red blood cell of ring-stage parasites increased the recovery rate and yielded a similar median number of genes per cell (red point) when compared to rings from un-lysed red blood cells.

**Fig. S2.**
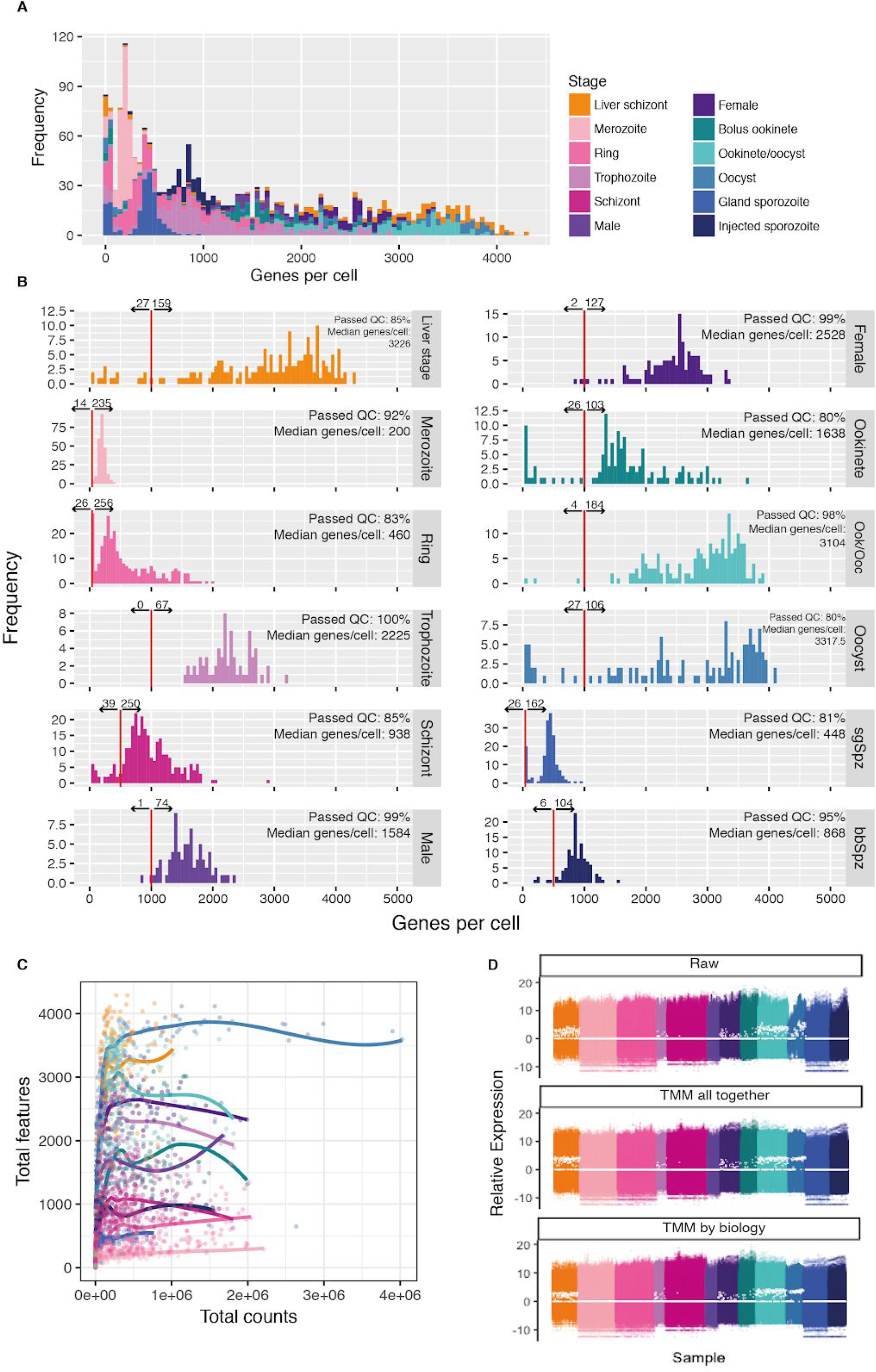
Quality control and normalization. **(A)** The number of genes detected per cell was dependent on life stage (*p*<0.001, ANOVA: Genes detected ~ parasite stage). **(B)** Poor quality cells were identified on a per stage basis based on the distribution of the number of genes detected per cell. The maximum number of genes per cell was found in oocysts, at day 4 post mosquito infection, with a median of 3318 genes per cell, while the minimum number of genes detected was found in merozoites with a median of 200 genes per cell. This is consistent with smaller cells having less total mRNA (*56*), and likely reflects biological differences in the mRNA content of each life stage. Interestingly, we detect almost twice the number of genes in sporozoites taken directly from injected saliva (n genes=868) relative to sporozoites that were isolated from salivary glands (n genes=448), suggesting a potential upregulation of gene expression at this transition. Quality control was also performed based on the number of reads per cell. The percentage value includes cells that were excluded based on genes and reads per cell. **(C)** Total counts per cell vs. total genes detected in all Smart-seq2 cells. Each stage is fit with a loess function. Sequencing saturation was likely reached because deeper sequencing did not result in an increase in the number of genes detected, irrespective of life stage. **(D)** Relative Expression plots for all cells across all genes. The top panel shows the relative expression of the raw counts data. The second panel shows the relative expression whereby cells have been TMM-normalised as one set. The bottom panel shows the same plot but for cells that have been subsetted into their respective groups of stages: Liver schizonts, IDC, gametocytes, ookinetes/oocysts, sporozoites, normalised by TMM, and then re-pooled. Based on the plots, normalisation method does not have a large effect on the relative expression; cell expression patterns have been smoothed in both methods. Unless otherwise stated, the grouped normalised data were used for further analyses.

**Fig. S3.**
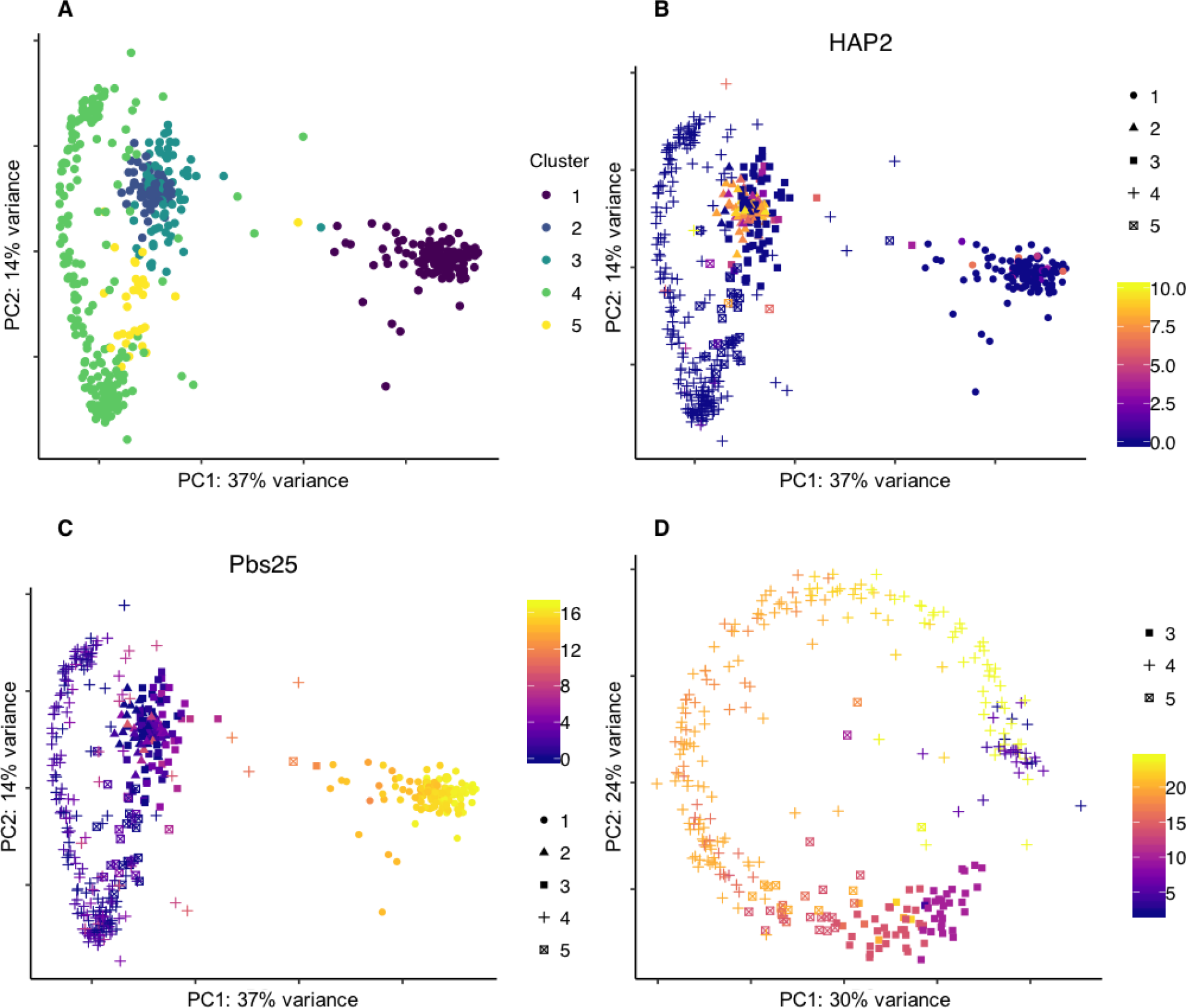
Classification of blood-stage parasites. Late-stage blood-stage parasites (trophozoites, schizonts and gametocytes) were purified from an overnight (20 h) culture of infected blood. Classification of these parasites was determined using single-cell consensus clustering in SC3 (*24*). (**A**) PCA of all late-stage parasites coloured by SC3 cluster assignment. (**B**) The PCA coloured by expression of HAP2, an established male marker gene (*29*), supporting assignment of cluster 2 cells as male gametocytes. (**C**) The PCA coloured by expression of Pbs25, an established female marker gene (*55*), supporting assignment of cluster 1 cells as female gametocytes. (**D**) PCA of asexual parasites (primarily trophozoites and schizonts). Points are shaped by SC3 cluster and coloured by the Spearman correlation with bulk time-course data from (*21*). A good correspondence between SC3 clusters and the bulk data is observed with trophozoites classified as 8-16 h, and schizonts greater than 16 h. A few parasites that cluster with schizonts had the strongest correlation with ring stages, which could be a potential contamination of a small number of rings in the purification process.

**Fig. S4.**
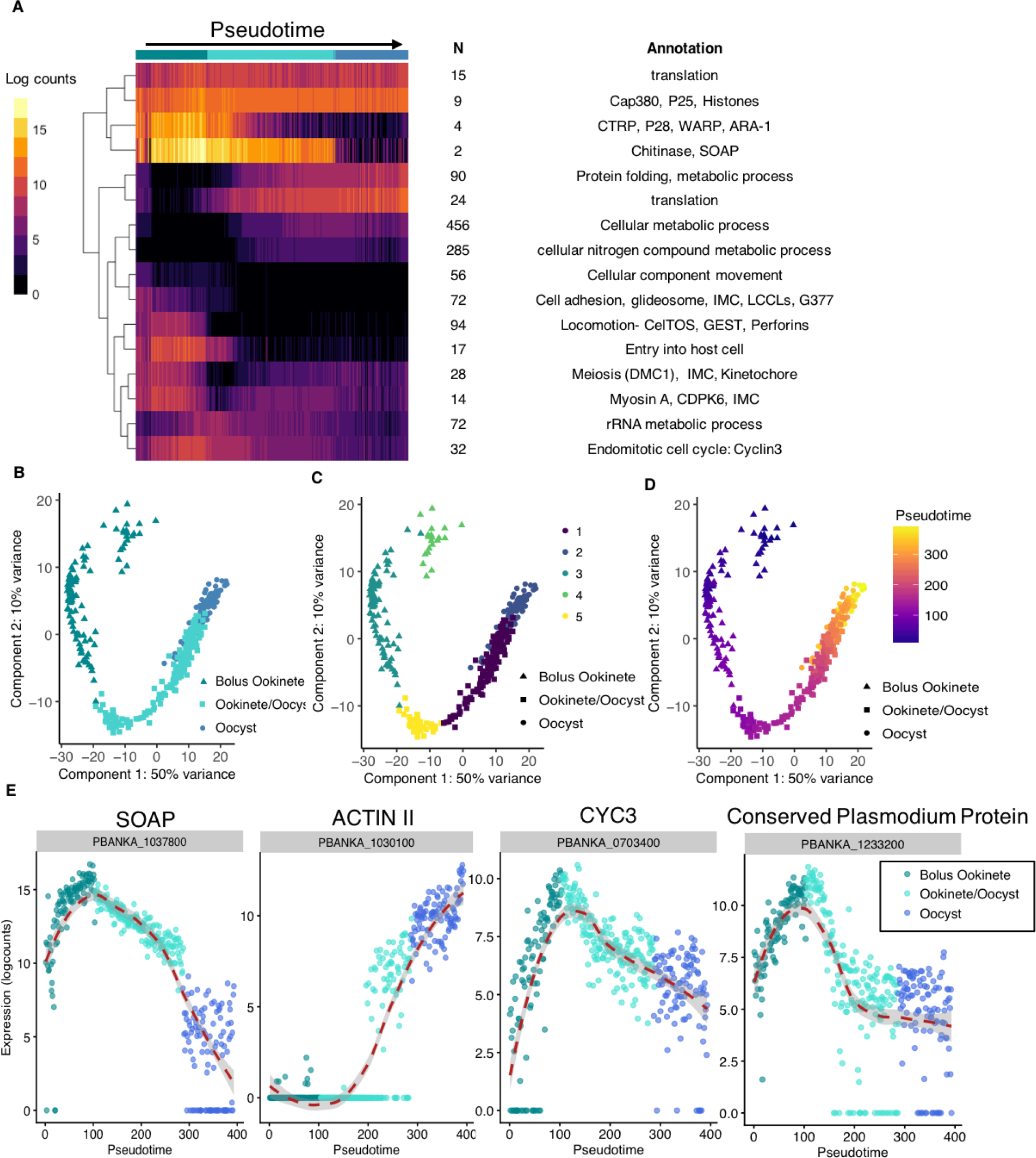
The ookinete to oocyst transition. In order to understand fine-scale changes in transcription over developmental time, we ordered cells from three experimental time-points (bolus ookinetes, the ookinete/oocyst transition period, and early oocysts) in pseudotime using SLICER (*39*). We identified 1270 genes that were differentially expressed over pseudotime among the 393 cells included. (**A**) A heatmap of the mean expression of each cluster of differentially expressed genes over pseudotime with manual annotation shows fine-scale patterns of expression over development. (**B**) PCA of stages represented by these cells. (**C**) The PCA coloured by their cluster assignments from SC3. Clusters three, four and five from were classified as ookinetes in further analysis, while clusters one and two were classified as oocysts based on expression of known marker genes. (**D**) The pseudotime ordering overlaid on the PCA. The pseudotime ordering matched both the different collection groups and the SC3 clusters. (**E**) Expression of four genes over pseudotime. SOAP is an ookinete marker genes (*54*). Actin II is an oocyst marker gene (*57*). We found other genes such as cyclin-3 (*58*) and PBANKA_1233200 expressed most highly at the transition between ookinete and oocyst stages. Cyclin-3 is known to be essential for oocyst development and we hypothesize that genes with a similar pattern of expression such as PBANKA_1233200 may be essential for the ookinete to oocyst transition.

**Fig. S5.**
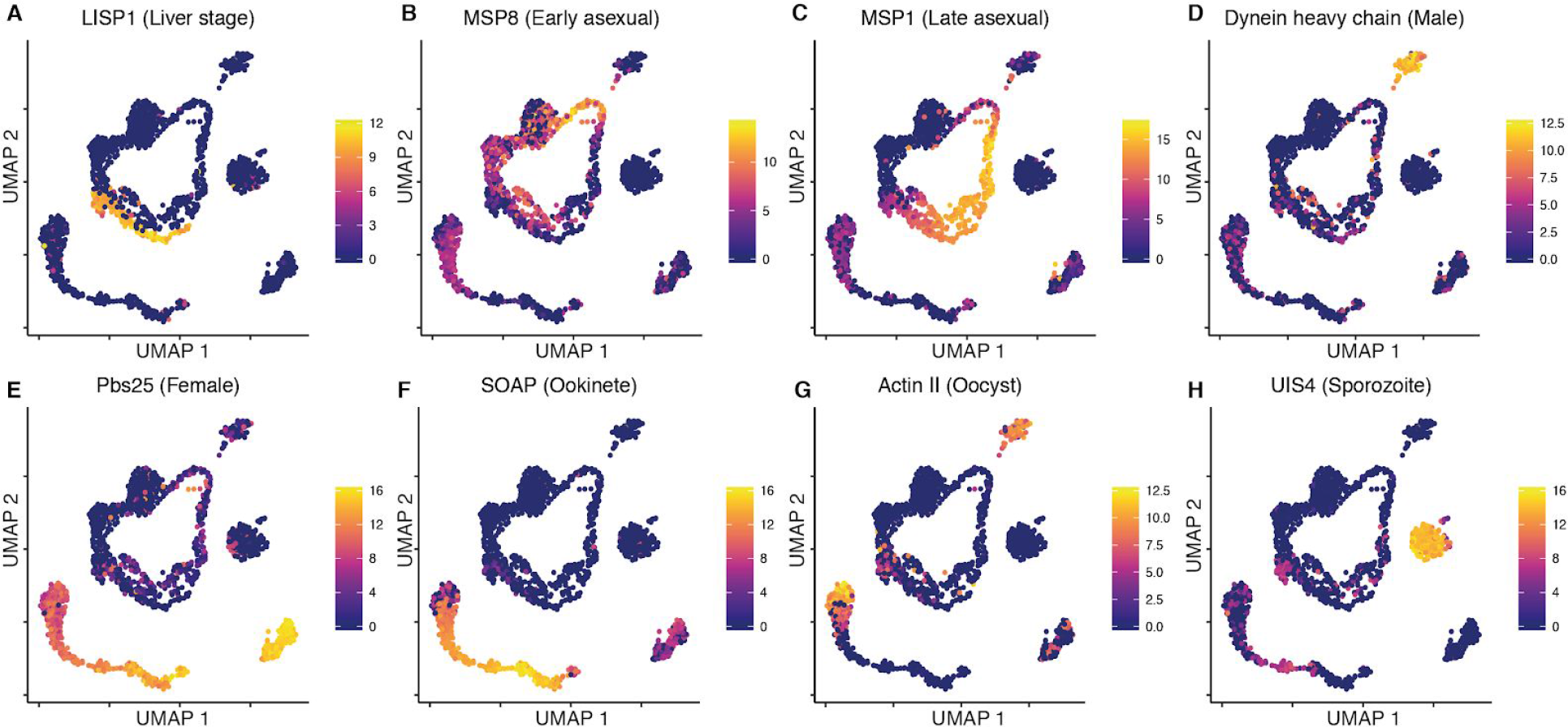
Marker gene expression. Expression of known marker genes corresponded with isolation method and stage classification. **(A)** LISP1 (*59*) is highly expressed in liver stages. (**B)** MSP8 (*60*) is highly expressed in early blood stage asexual parasites. **(C)** MSP1 (*61*) is highly expressed in late blood stage asexual parasites. **(D)** Dynein heavy chain (PBANKA_0416100) (*62*) is highly expressed in male gametocytes. **(E)** Pbs25 (*55*) is highly expressed in female gametocytes. (**F)** SOAP (*54*) is highly expressed in ookinetes. **(G)** Actin II (*57*) is highly expressed in oocysts. **(H)** UIS4 (*63*) was highly expressed in sporozoites.

**Fig. S6.**
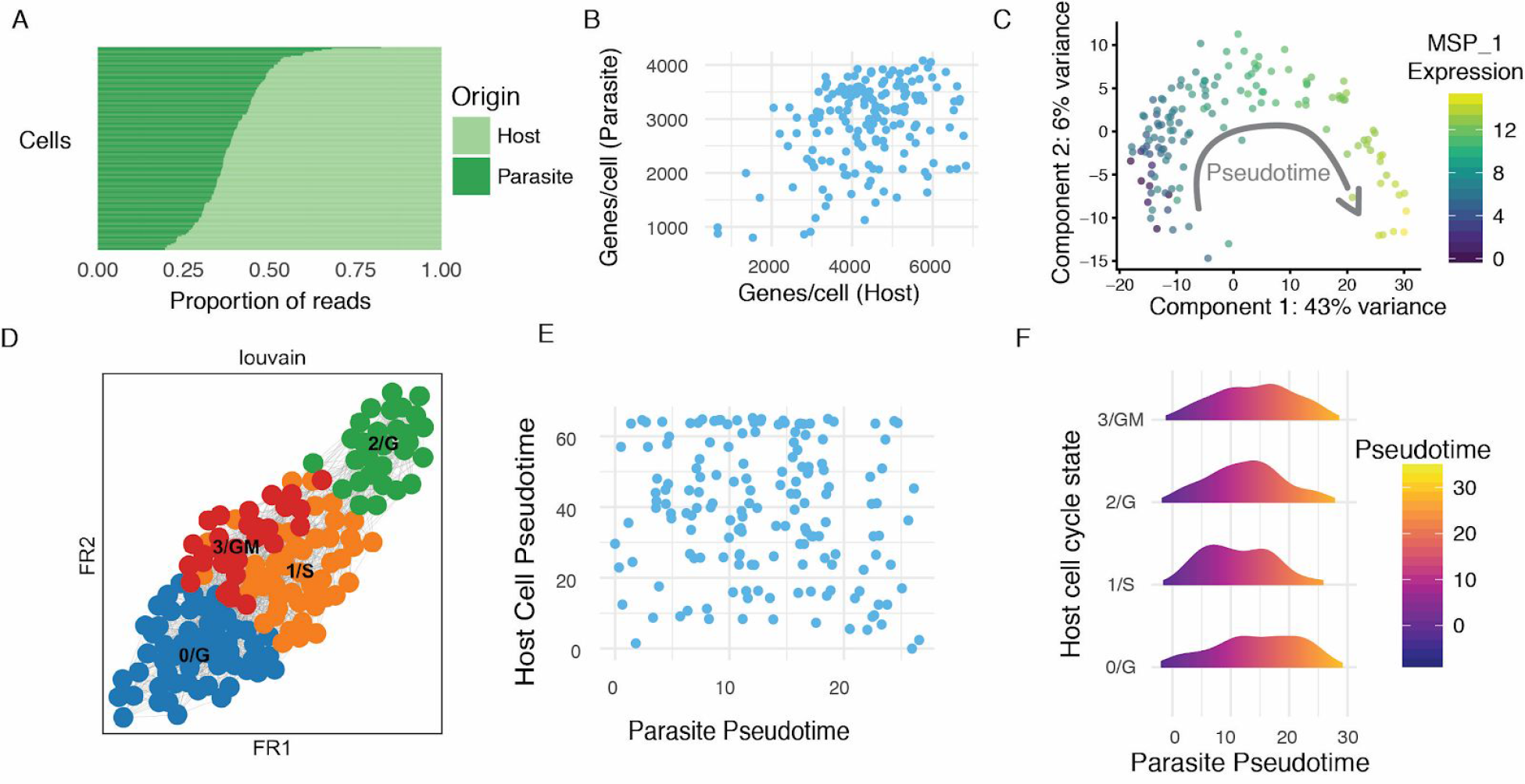
Dual scRNA-seq of host and parasite transcriptomes during exo-erythrocytic schizogony. Transcriptomes were generated from HeLa cells containing mCherry liver stage parasites 44 h post infection and selected based on fluorescence **(A)** Proportions of reads mapping to either the host (*H. sapiens*) or the parasite (*P. berghei*) genomes. **(B)** Number of genes per cell identified in host and parasite transcriptomes. **(C)** PCA of parasite transcriptomes identifies a developmental progression of the liver stages that corresponds to a known marker of progressing exo-erythrocytic schizogony (MSP-1, PBANKA_0831000 (*64*)). **(D)** Force directed graph (*65*) of host transcriptomes, with different louvain clusters identified as different cell-cycle states. **(E)** Pseudotime analysis for both host and parasite transcriptomes were computed using SLICER (*39*) and plotted against one another showing no correlation of developmental state between the host cell and the parasite cell that resided in it. **(F)** Cell-cycle state of the host cell also did not correspond to a particular pseudotime of the parasite indicating a decoupling between host cell-cycle state and parasite developmental progression.

**Fig. S7.**
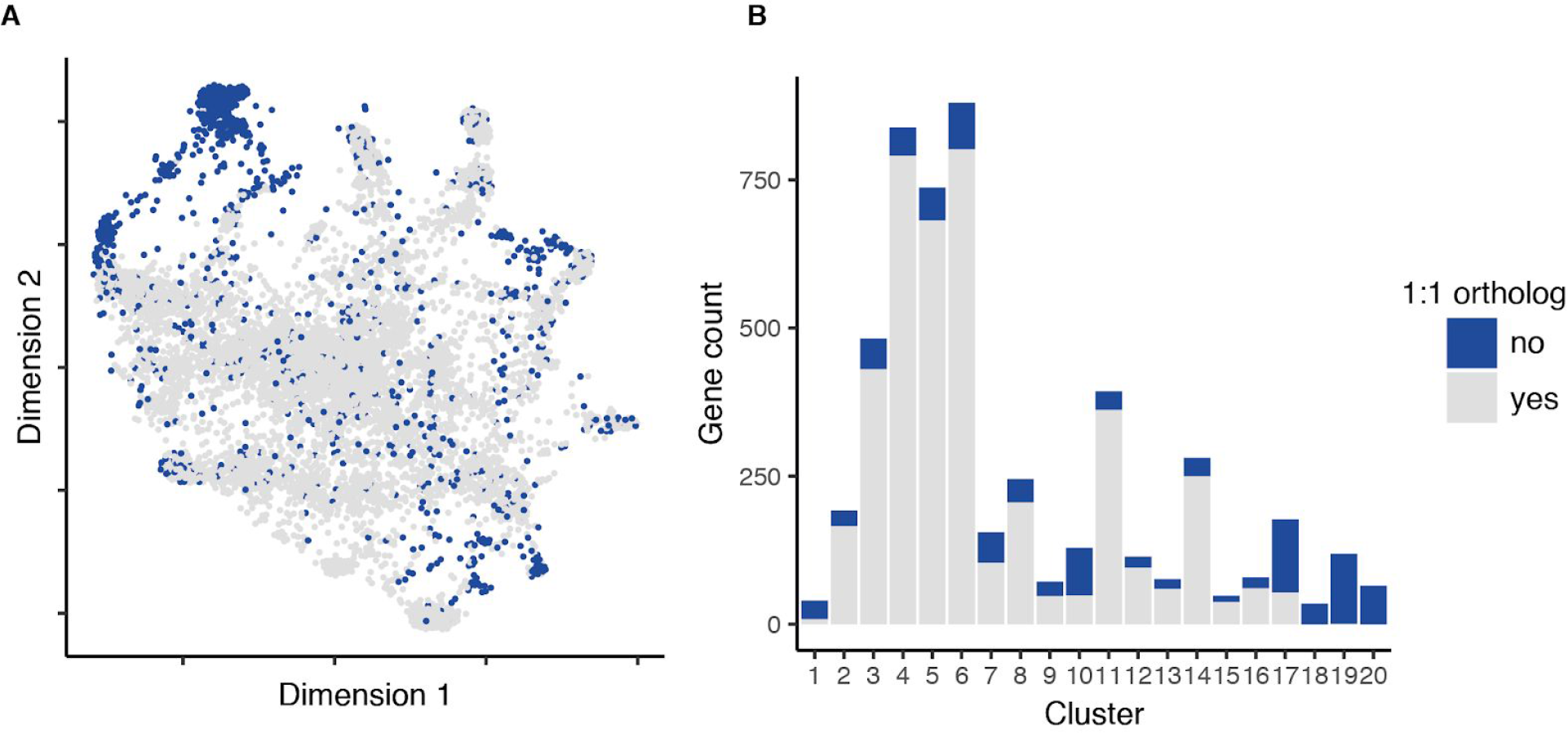
Conservation across kNN graph. **(A)** The kNN graph with non-orthologous (*P. berghei* to *P. falciparum*) genes highlighted in blue. **(B)** A barplot of the gene counts for each cluster colored by orthology. Clusters 1, 7, 10, 17-20 were significantly enriched for genes that have no ortholog with *P. falciparum* (Fisher’s exact test, bonferroni *p* <0.001). Clusters 18, 19, and 20 were composed almost completely of genes from multigene families containing no orthologs in *P. falciparu*m.

**Fig. S8.**
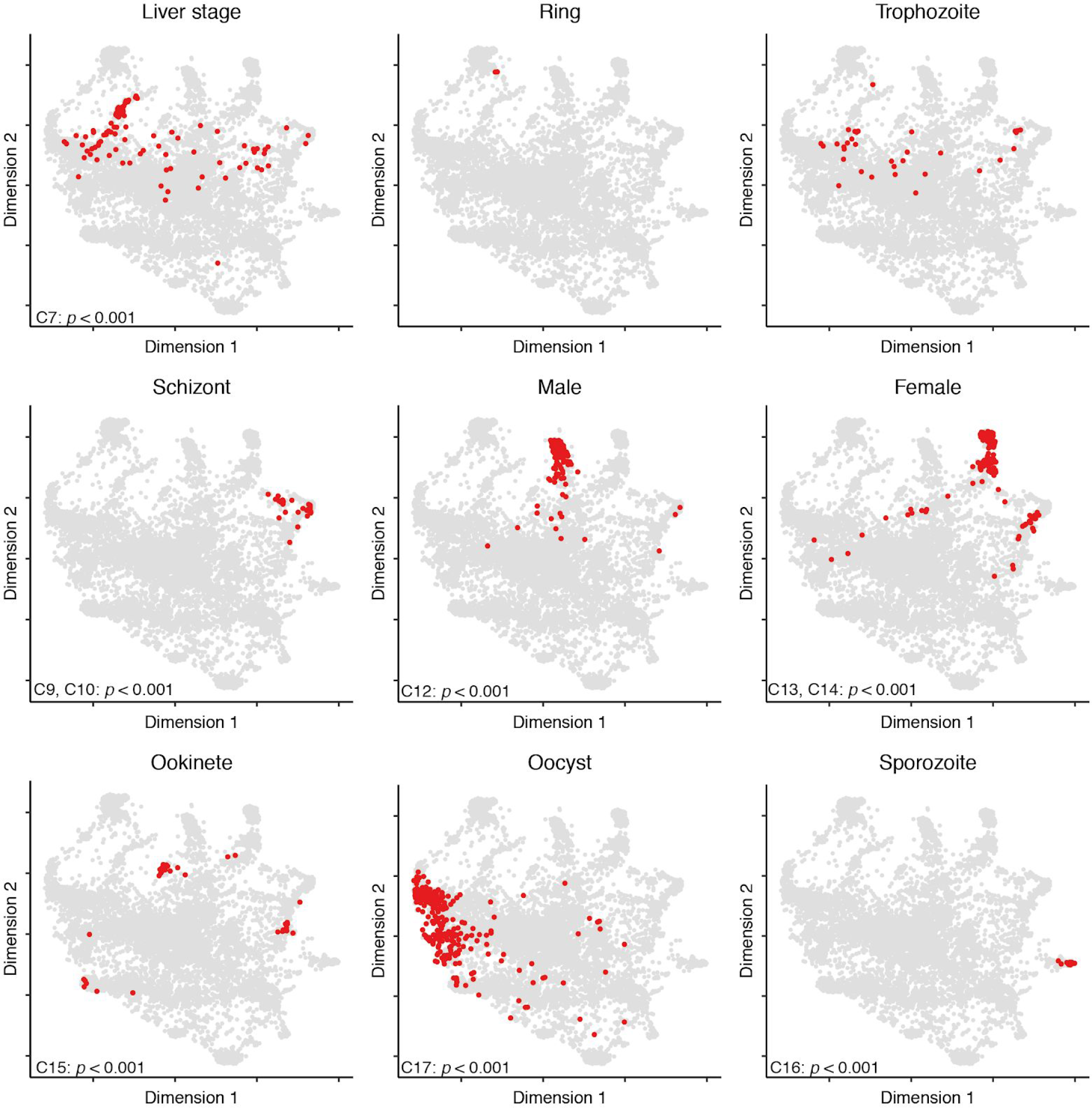
Unique core expression genes by stage across the kNN graph. For each stage, a unique core set of genes fe was defined by identifying the genes for each stage expressed in more than fifty percent of 60 randomly selected cells from that stage. Core genes unique to that stage were considered the ‘unique core’. These genes are highlighted for each stage in red on the kNN graph. We see a clear pattern of clustering for most stages (see inset p-values) of unique core genes that matches the graph spectral clustering shown in Fig. 2. Enrichment of core unique genes was calculated for each cluster using a Fisher’s exact test.

**Fig. S9.**
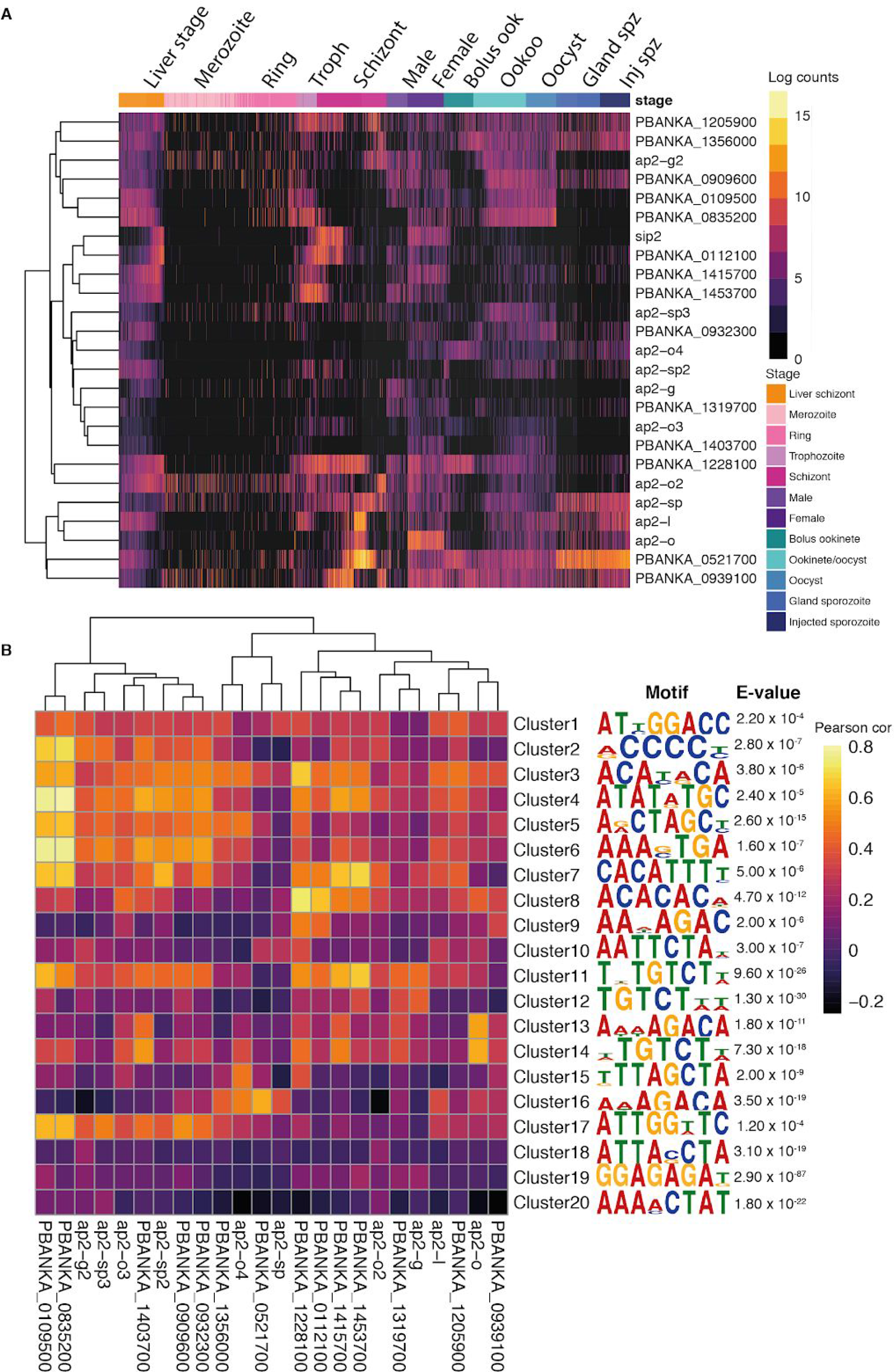
Expression of AP2 transcription factors across all cells in the data set. **(A)** A heatmap of log expression counts of 25 AP2 transcription factors across all cells in the data set. Cells are ordered in a developmental trajectory based on life cycle order and pseudotime. **(B)** Heatmap of Pearson correlations between AP2 gene expression and gene clusters from Fig. 2. The column on the right shows the most significantly enriched motif in that cluster with the corresponding e-value (see also file S1). We found that the most significant motif of cluster 15, which is highly expressed in the ookinete stage, matched the AP2-O motif ([G/C]TTAGCTA in our analysis, [TC][AG]GC[TC][AG] in (*66*)). The AP2-O TF is essential for ookinete development and the gene is highly expressed in the preceding female stage (*66*). This validation suggests that our analysis can identify as-yet unknown TF binding motifs in *Plasmodium*.

**Fig. S10.**
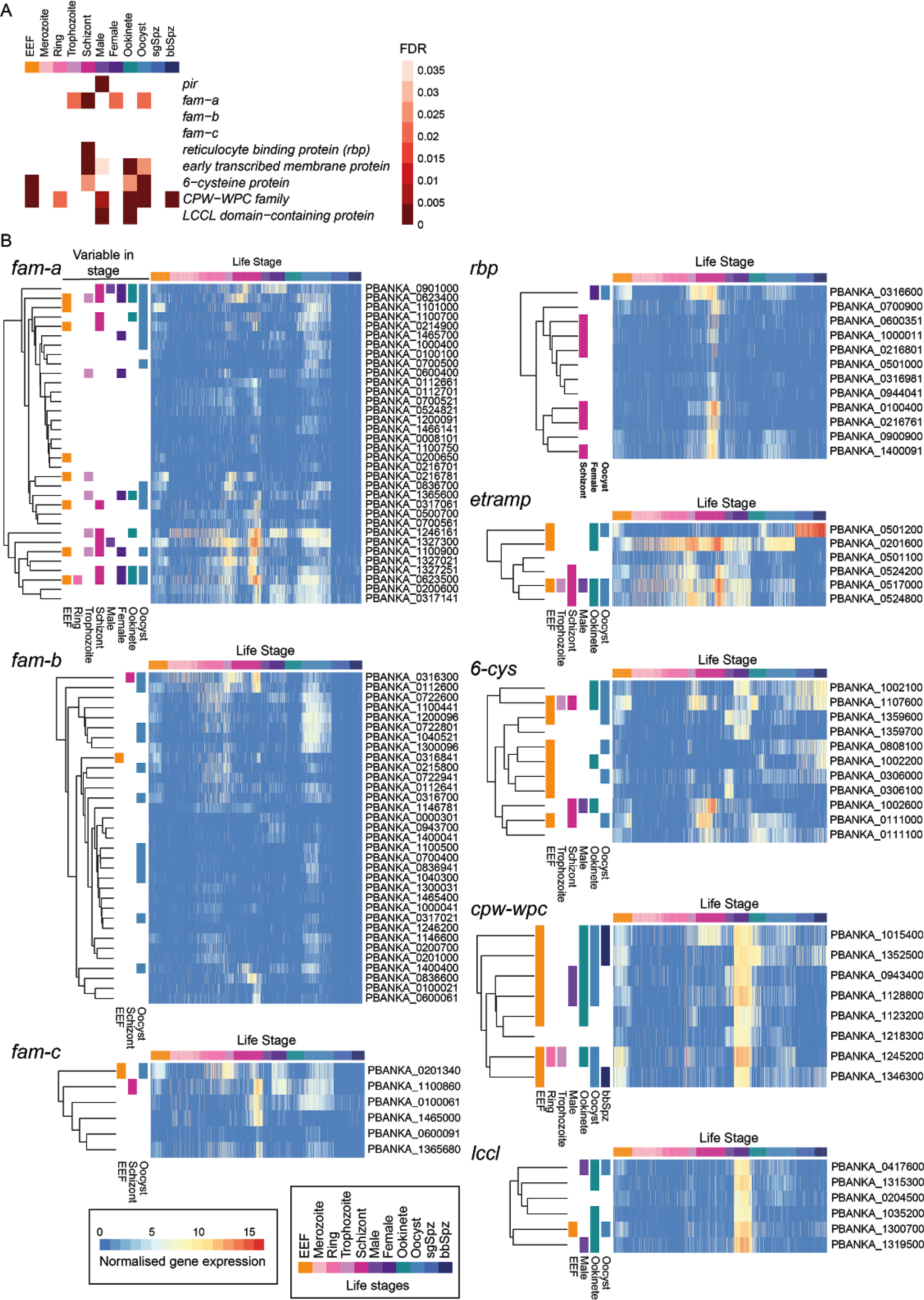
Multigene families show development-independent variable expression between cells in most life stages. **(A)** A heatmap shows which stages were enriched for variability in expression of each multigene family. Variable genes were identified in each stage using Smart-seq2 data and controlling for development by regressing out pseudotime in liver stage EEFs, merozoites, rings, trophozoites, schizonts, ookinetes and oocysts. The hypergeometric test was used to determine enrichment of each gene family amongst variable genes in each stage and the resulting *p*-values were adjusted using the False Discovery Rate (FDR). **(B)** Each heatmap shows the TMM-normalised expression levels of members of a gene family over the life cycle of *P. berghei*. The data were filtered to show only those genes with at least 10 read counts in at least 10 cells. Genes were clustered based on their expression pattern. Genes found to vary in expression level between cells of the same stage independent of development are highlighted to the left of each heatmap. The presence of a turquoise square, for example, indicates that that gene was variably expressed between ookinetes.

**Fig. S11.**
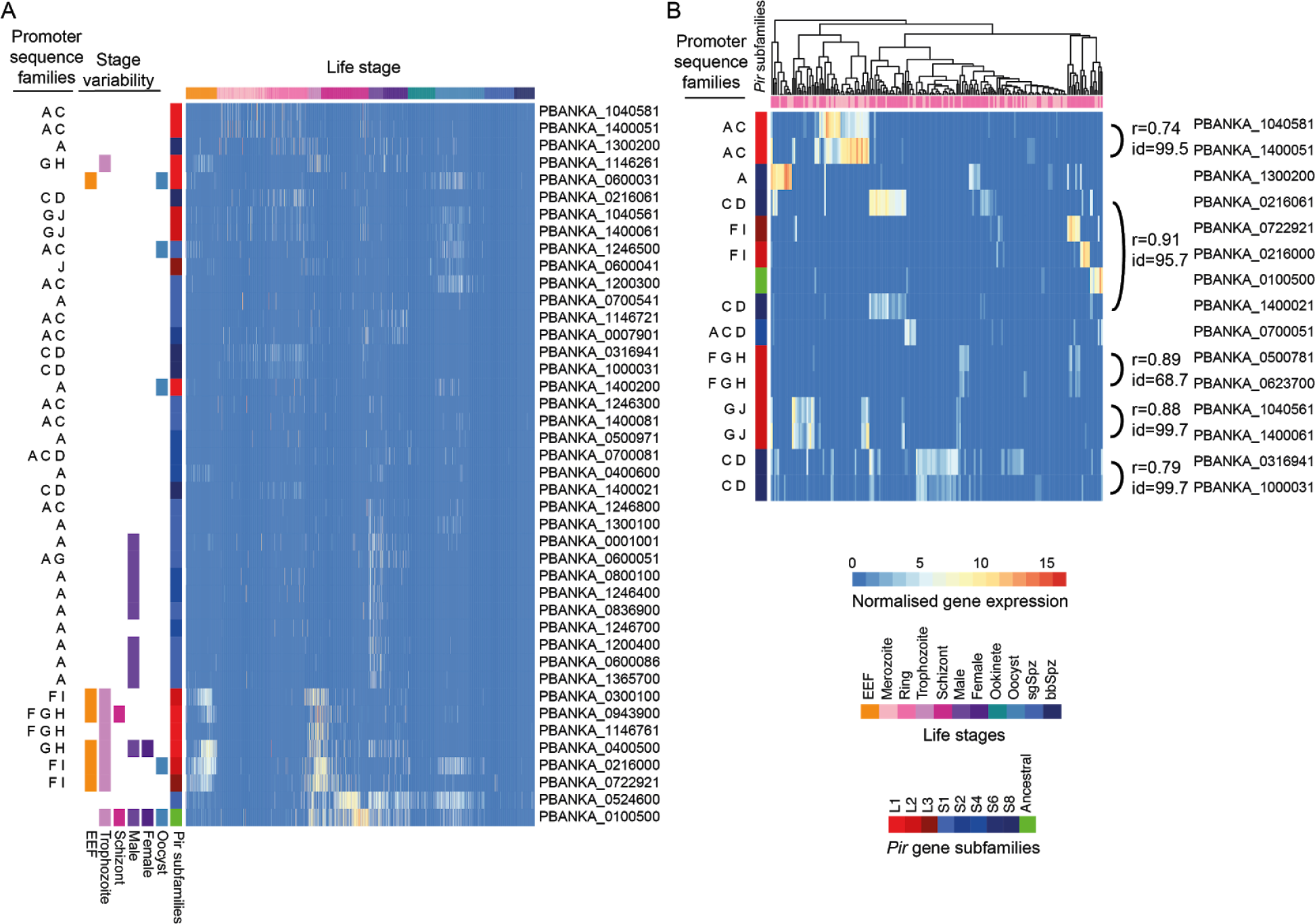
The *pir* gene family shows distinct patterns of expression and promoter architecture in several parts of the life cycle. **(A)** The heatmap shows the TMM-normalised expression levels of members of the *pir* gene family over the life cycle of *P. berghei*. The data were filtered to show only those genes expressed with at least 10 read counts in at least 10 cells over the whole dataset. Genes were clustered based on their expression pattern. The presence of an orange square, for example, indicates that that gene was variably expressed between liver stage EEFs. The subfamily classification of each *pir* gene is indicated in red colours (L-type *pir* genes), blue colours (S-type *pir* genes) and green (ancestral *pir* gene) to the left of the heatmap. Families of sequence identified in the promoter regions of each gene are indicated with letters A-J. One cluster of *pir* genes variably expressed in both EEFs and trophozoites tended to have ‘F’ sequences in their promoters. Another cluster was variably expressed in male gametocytes and tended to have promoters with ‘A’ sequences. **(B)** This heatmap shows merozoites and rings and *pir* genes expressed with at least 10 read counts in at least 10 of these cells. Both genes and cells were clustered based on their expression levels. Five pairs of genes show evidence of co-expression (Pearson r > 0.7). Each pair tended to have high sequence identity (BLAST sequence identity > 95%), the same promoter architecture (identical pattern of upstream sequence families) with each member of a pair residing on different chromosomes.

**Fig. S12.**
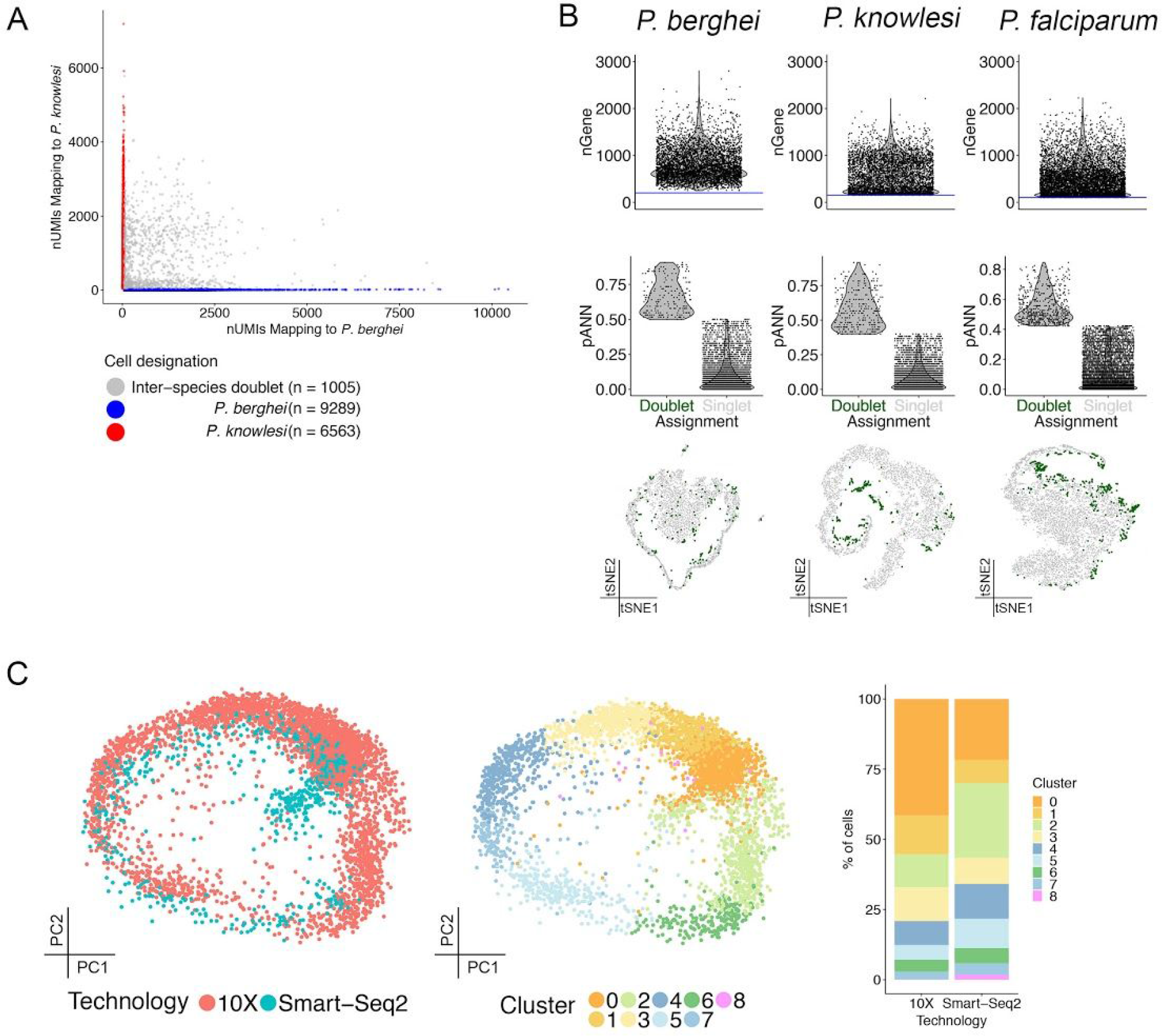
Comparison of Smart-seq2 data with 10X data and technical assessment. **(A)** A ‘barnyard’ plot showing the doublet rate between the *P. knowlesi* and *P. berghei* mixed species experiment. **(B)** For each species, Top: a violin plot showing the number of genes detected per cell (nGene) with a line representing the QC threshold used; Middle: A violin plot showing the proportion of artificial nearest neighbors (pANN) as calculated by DoubletFinder (*38*), with cells split by their final assignment; Bottom: A tSNE plot showing doublets highlighted in green against singlets in grey, showing the distribution of doublets among stages. **(C)** PCA plot overlaying the Smart-seq2 to the 10X data using a CCA corrected distance matrix coloured by technology (left) and cluster (middle). The proportional representation of these clusters captured by each technology is displayed on the right.

**Fig. S13.**
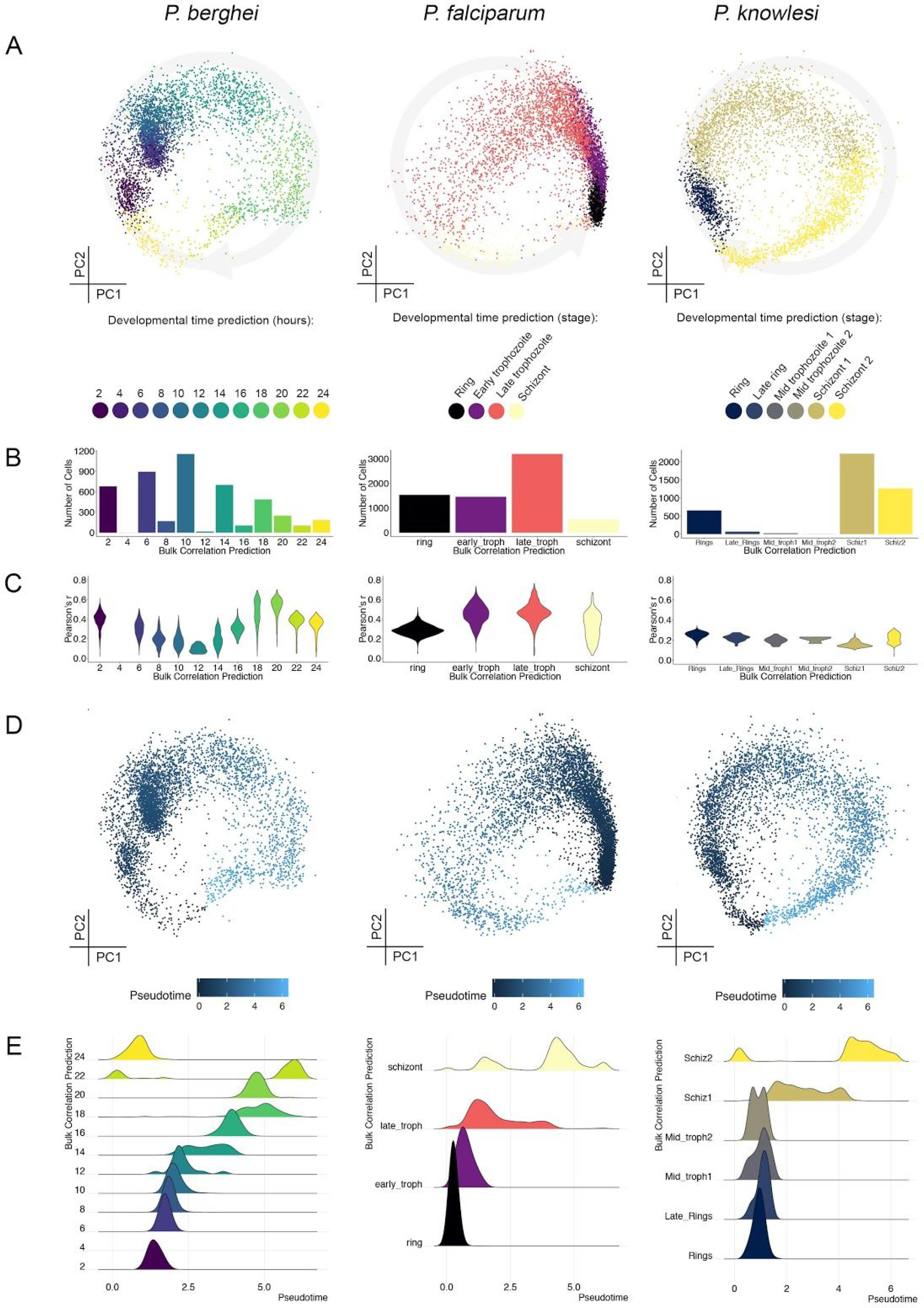
Correlation of 10X data sets with bulk expression data. **(A)** PCA plots for each species, coloured by their predicted life cycle stage according to maximum correlation with the following bulk time course reference data sets: *P. berghei* microarray (*21*); *P. falciparum* RNA-seq (*67*); *P. knowlesi* microarray (*68*). **(B)** Distribution of cells according to predicted life cycle stage based on bulk data. The temporal density of sampling over the IDC is different between bulk data sets so this results in a different number of bins for each species. **(C)** Violin plots showing the Pearson correlation coefficient, r, for each species by predicted life cycle stage. **(D)** PCA plots for each species, coloured by pseudotime value, calculated using the custom clock approach. **(E)** Density ridgeline plots showing the relationship between pseudotime and predicted life cycle stage.

**Fig. S14.**
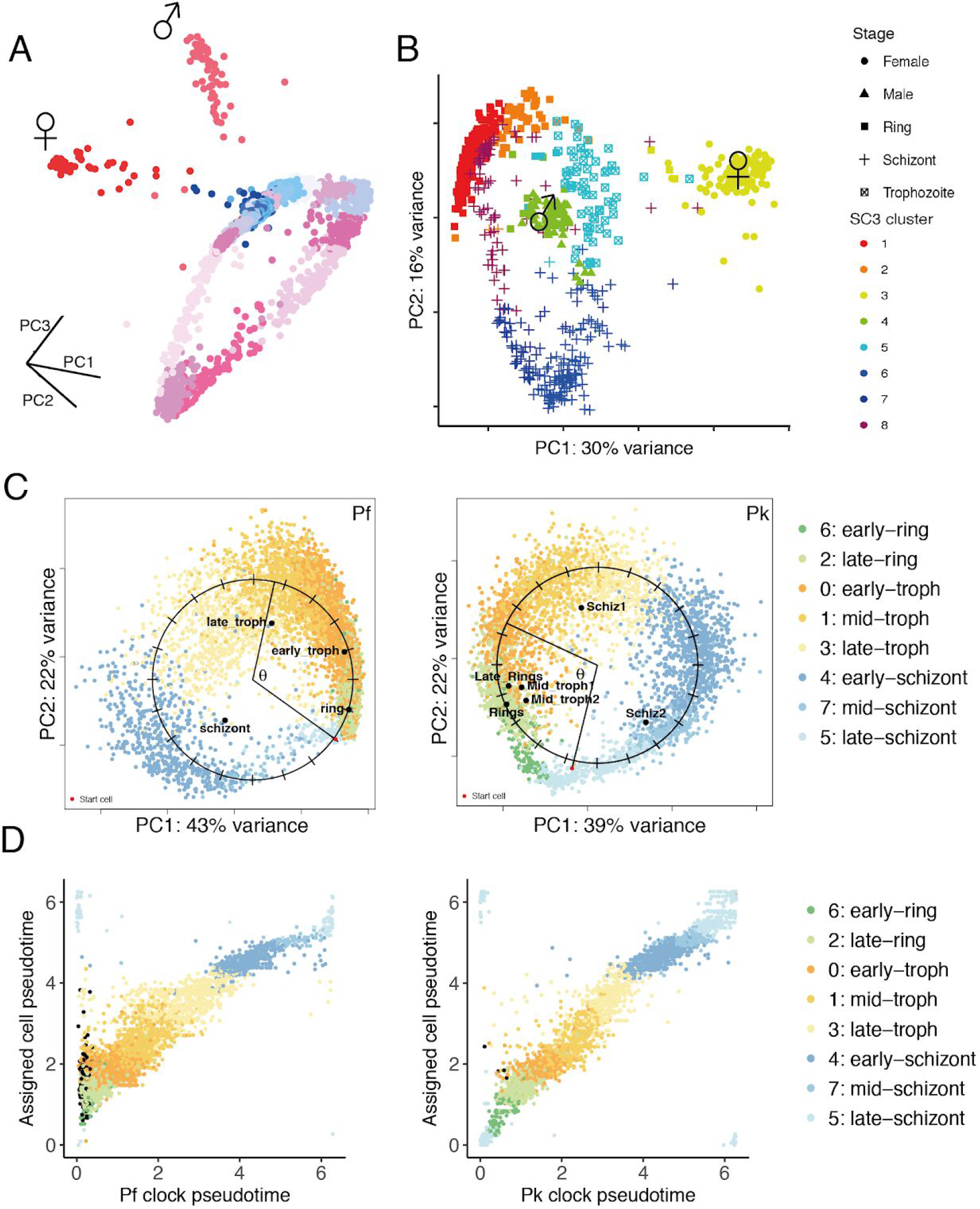
SCmap of 10X data. **(A)**In order to identify male and female gametocytes, clustering of 10X data was initially done with the Seurat shared nearest neighbor modularity optimization method, which identified 26 clusters, two of which corresponded to mature male and female gametocytes based on known markers. Cells are shown on a PCA and colored by cluster assignment. **(B)** SC3 clustering of blood-stage parasite data (rings, trophozoites, schizonts, and mature gametocytes). Cells were re-clustered in SC3 into eight clusters that were then used for the cluster assignment shown in Fig. 3A. **(C)** ‘Clock’ pseudotime was calculated independently for both *P. falciparum* (left) and *P. knowlesi* (right) and the mean coordinates of the bulk prediction were mapped onto the PCA (black points). **(D)** The correspondence between the independent pseudotime calculation from and the pseudotime of the assigned *P. berghei* cell based on scmap There was a strong correlation between pseudotime for each species and the assigned cell pseudotime further supporting the cell assignments from scmap.

**Fig. S15.**
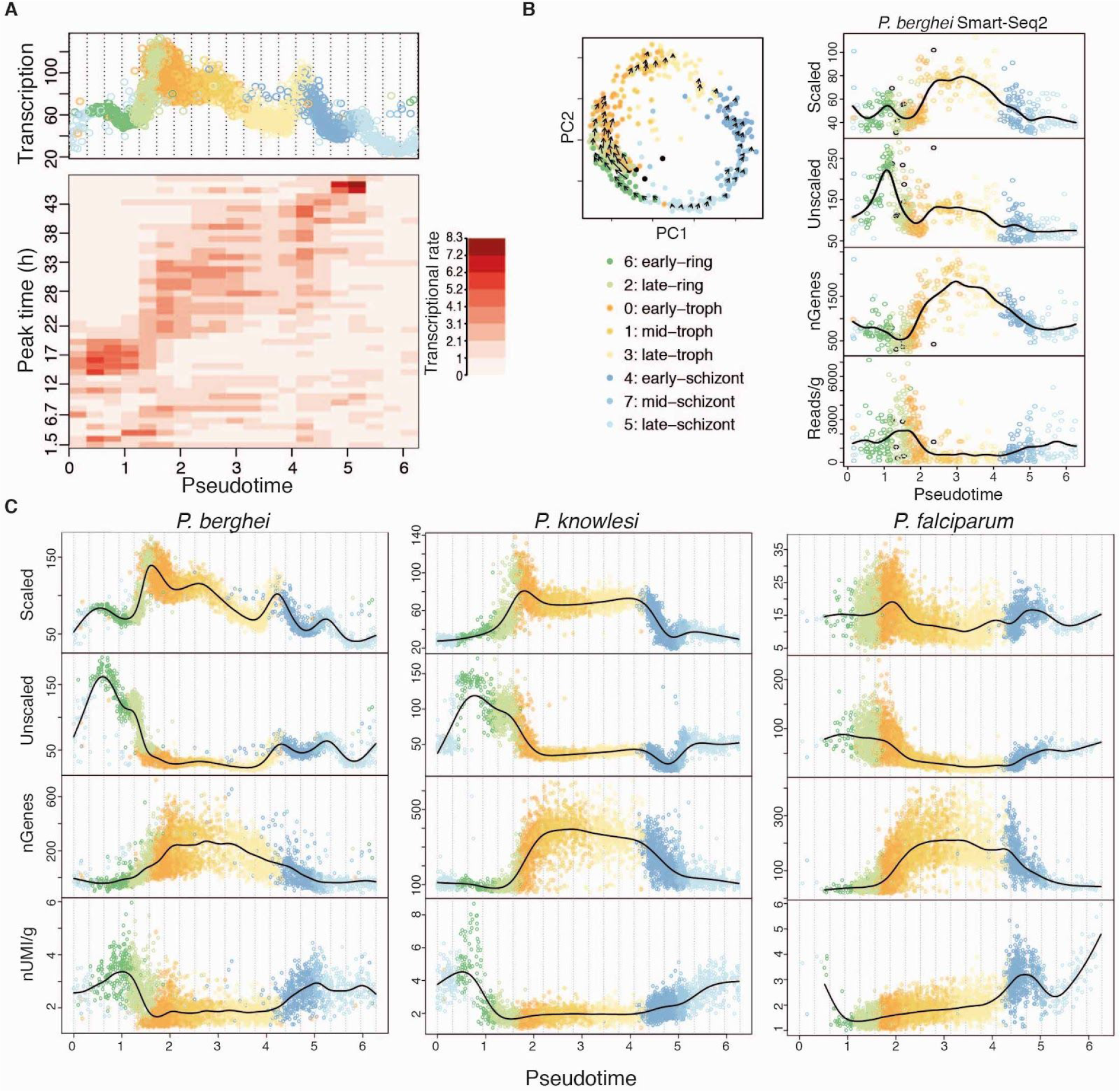
Transcriptional rate varies across the IDC. **(A)** A heatmap showing the average scaled transcriptional rate of genes in *P. berghei* as measured by RNA velocity (*19*) in the IDC. Genes are ordered and binned along the vertical axis by their peak time in *P. falciparum* in (*20*). The top panel shows the scaled transcriptional rate of each cell over pseudotime in the *P. berghei* 10X data. The groups of genes from (*20*) also had high transcriptional rates at corresponding points in the life cycle in *P. berghei* based on RNA velocity. **(B)** RNA velocity of the Smart-seq2 IDC cells. Cells were mapped to the 10X *P. berghei* reference index with scmap-cell to assign each cell a cluster and a pseudotime value allowing us to directly compare the two data sets. Cells are colored by their assigned cell cluster. The left plot is a PCA of the 548 Smart-seq2 IDC cells with the arrows representing the local average velocity. The right panel displays the same cells over pseudotime showing the relative increase in transcription across all transcripts unscaled or scaled by gene, followed by the number of genes detected and the number of reads per gene in each cell over pseudotime. **(C)** The three species 10X data sets over assigned cell pseudotime based on the *P. berghei* 10X reference index. Cells are colored by their assigned cell cluster. Each panel shows the relative increase in transcription (scaled and unscaled) as well the the number of genes detected and the number of UMIs per gene. We observed a similar pattern of transcriptional dynamics in both the Smart-seq2 and 10X *P. berghei* data. Additionally, the pattern of early stages having the highest transcriptional rate was seen across species, suggesting changes in transcriptional rates over the IDC are highly conserved.

**Fig. S16.**
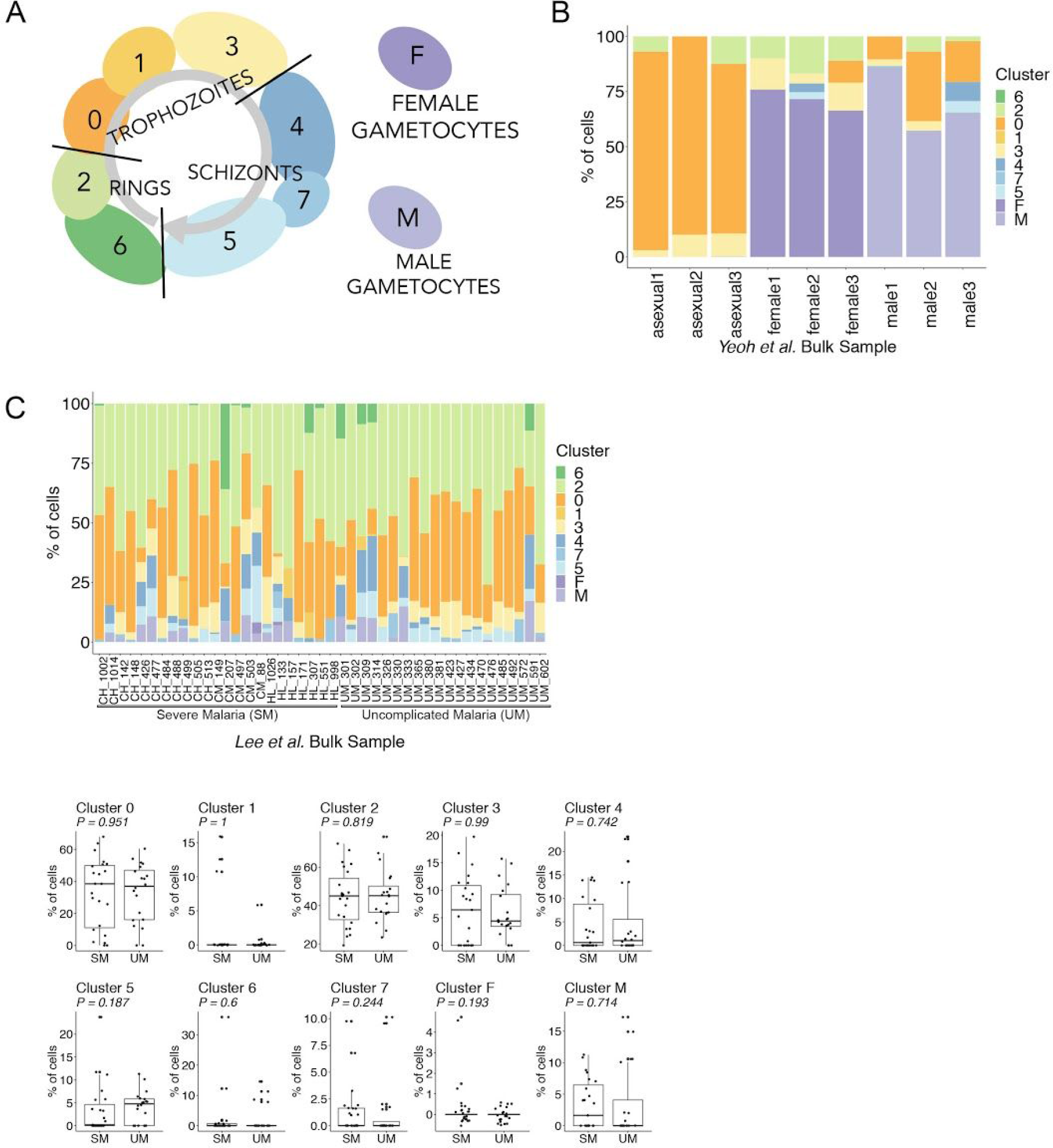
Deconvolution of bulk transcriptomic samples using scRNA-seq data. **(A)** A diagram of the *P. berghei* 10X data from fig. S14, which is used as a reference to call markers for each cell cluster. **(B)** The *Yeoh et al.* study (*22*) used the 820cl1m1cl1 *P. berghei* parasite line (*69*), which has RFP females and GFP males, to examine each sex separately by RNA-seq. Asexual samples were obtained straight from a mouse and only circulating forms are expected to be present. Using BSeq-SC (*53*) to estimate cell type proportions present in the bulk data, only circulating forms are observed in the asexual samples, as well as a majority of females in the female samples, and a majority of males in the male samples. Additional cells called in the gametocyte samples are hypothesised to be developing gametocytes, technical noise in our method, and/or true asexual cells that are present in the sorting gate. **(C)** We tested whether the *P. berghei* cell atlas could be used to deconvolve bulk RNA-seq data from *P. falciparum* samples collected from patients with uncomplicated or severe malaria (*23*). We find that the cell population in the bulk data is composed primarily of early stage circulating forms as expected (top). We also observe cell type composition is independent of the severity of malaria (bottom) as described by the authors in this study. The Malaria Cell Atlas provides a robust index by which to deconvolve bulk transcriptomic data into contributing cell populations.

**Fig. S17.**
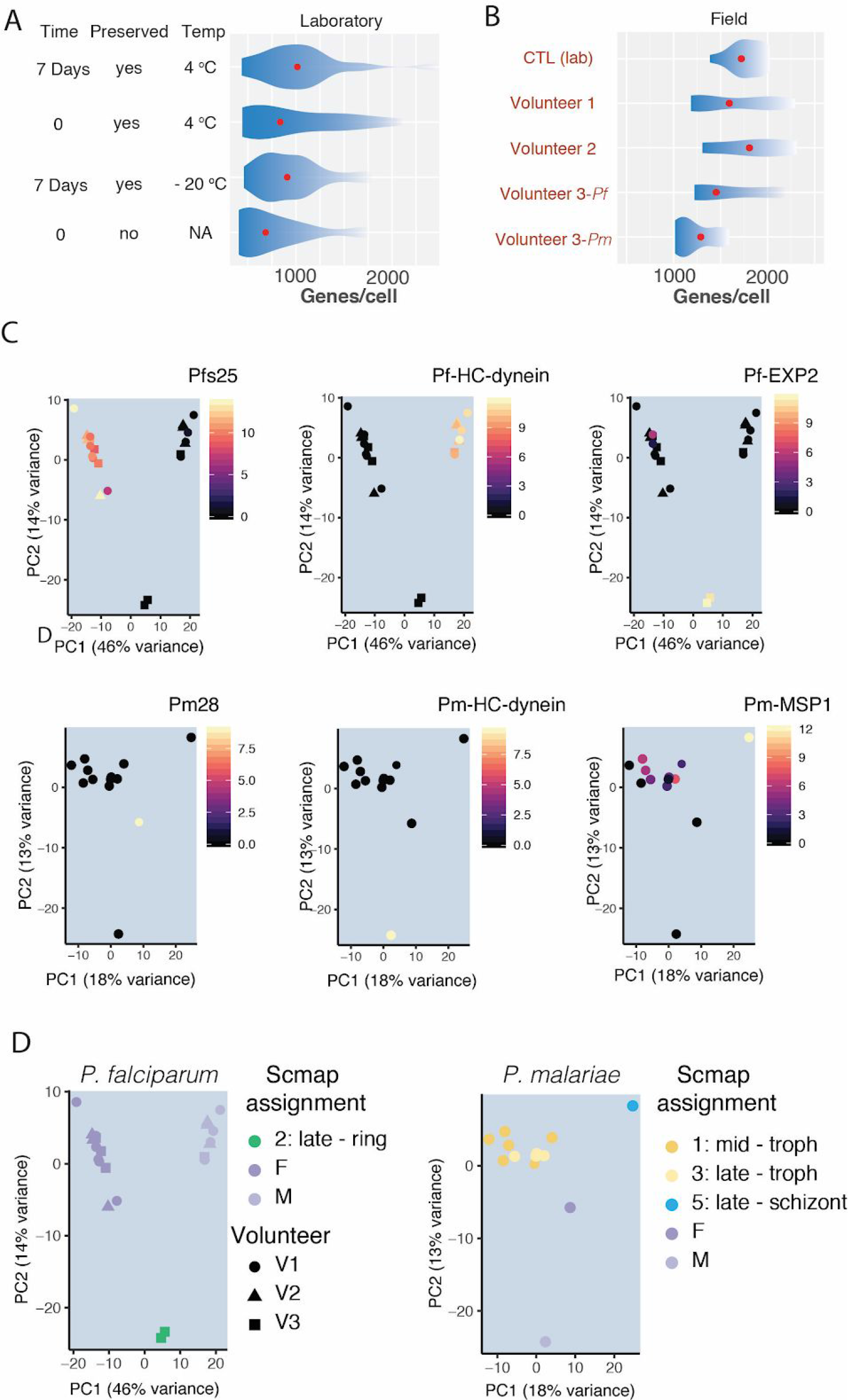
Preservation and analysis of single cell transcriptomes from natural carriers of *Plasmodium*. **(A)** *P. falciparum* trophozoites and schizonts were preserved in 80% methanol and kept at different temperatures for varying amounts of time before analysis by Smart-seq2; this revealed that transcriptomes from methanol fixed parasites are of equivalent quality than ones from RNALater fixed parasites. **(B)** We used our preservation protocol on parasites collected from naturally-infected carriers in Mbita Kenya and established that even after three weeks of preservation, high quality single cell transcriptomes were possible to retrieve using Smart-seq2. **(C)** PCA of 22 *P. falciparum* and 13 *P. malariae* wild single cell transcriptomes showing expression of known (*P. falciparum* (*70*)) or putative (*P. malariae*) male (HC-dynein), female (Pfs 25 and Pm28), early ring (EXP2), and putative late stage markers (MSP-1) **(D)** PCAs of the same cells as in (C) but with their scmap assignment based on the *P. berghei* 10X Malaria Cell Atlas. Life stages can be assigned using known markers as demonstrated in (C) but to place cells in developmental time, the full atlas is required (see Fig. 4B for placement of these cells).

